# miR-329 and miR-495-mediated Prr7 downregulation is required for homeostatic synaptic depression in rat hippocampal neurons

**DOI:** 10.1101/2022.03.08.483397

**Authors:** Michiko O. Inouye, David Colameo, Irina Ammann, Gerhard Schratt

## Abstract

Homeostatic synaptic depression (HSD) in excitatory neurons is a cell autonomous mechanism which protects excitatory neurons from over-excitation as a consequence of chronic increases in network activity. In this process, excitatory synapses are weakened and eventually eliminated, as evidenced by a reduction in synaptic AMPA receptor expression and dendritic spine loss. Originally considered a global, cell-wide mechanism, local forms of regulation, such as the local control of mRNA translation in dendrites, are being increasingly recognized in HSD. Yet, identification of excitatory proteins whose local regulation is required for HSD is still limited. Here, we show that Proline-rich protein 7/Transmembrane Adapter Protein 3 (Prr7) downregulation in dendrites of rat hippocampal neurons is necessary for HSD induced by chronic increase in network activity resulting from a blockade of inhibitory synaptic transmission by picrotoxin (PTX). We further identify two activity-regulated miRNAs, miR-329-3p and miR-495-3p, which inhibit Prr7 mRNA translation and are required for HSD. Moreover, we found that Prr7 knockdown reduces expression of the synaptic scaffolding protein SPAR, which is rescued by pharmacological inhibition of CDK5, indicating a role of Prr7 protein in the maintenance of excitatory synapses via protection of SPAR from degradation. Together, our findings highlight a novel HSD mechanism in which chronic activity leads to miR-329 and miR-495-mediated local Prr7 reduction upstream of the CDK5-SPAR pathway.

## Introduction

Homeostatic synaptic depression (HSD) is a type of homeostatic plasticity by which excitatory neurons compensate for increased network activity to maintain a physiological range of excitatory transmission (reviewed in Turrigiano, 2008; Yu & Goda, 2009; Turrigiano, 2012). Adaptive mechanisms to maintain neuronal homeostasis include changes in synaptic AMPA receptors (Seeburg et al., 2008), and spine number (Kirov et al., 1999: Wierenga et al., 2006). Abnormal dendritic spine density and altered AMPAR internalization have been suggested in epilepsy (Isokawa et al., 1997), schizophrenia (Glantz et al., 2000), and autism spectrum disorder (Hutsler and Zhang, 2010), as well as in disease models of Fragile X (Fmr1-knockout) (Jawaid et al., 2018) and Rett Syndrome (Mecp2-mutant) (Chao et al., 2007), highlighting the importance of HSD regulation for neuronal homeostasis.

Although the majority of studies on HSD have utilized GABA-receptor antagonists (e.g. Picrotoxin (PTX), Bicuculline) to investigate HSD in response to network-wide stimulation, a study employing optogenetic methods of stimulation demonstrated that the mechanism of HSD is cell-autonomous (Goold and Nicoll, 2010). Namely, upon 24h photostimulation of channel rhodopsin-2 (ChR2)-expressing CA1 pyramidal neurons, lowered mEPSC frequency and dendritic spine number were observed, indicating that individual neurons possess intrinsic mechanisms to regulate their synapse number in response to chronic activity. The spine loss in HSD is supported by a number of other studies using either optogenetics (Mendez et al., 2018) or pharmacological stimulation (Pak and Sheng, 2003; Fiore et al., 2014; Chowdhury et al., 2018).

Pathways underlying HSD have been examined in detail. In a well-studied mechanism, elevated synaptic activity first causes L-type voltage-gated calcium channel opening and NMDAR activation, which results in calcium influx (Goold and Nicoll, 2010, Pak and Sheng 2003). Calcium binds calmodulin, which initiates a cascade of CaM kinases (Wayman et al., 2008), of which CaMKK and CaMK4 are required for HSD (Goold and Nicoll, 2010). The cascade transcriptionally activates Polo-like kinase 2 (Plk2/SNK), a member of the polo family of serine/threonine protein kinases, over a time scale of hours (Kauselmann et al., 1999; Pak and Sheng, 2003). The induced Plk2 is targeted to dendritic spines and binds to a PSD-95 interacting factor, spine-associated Rap guanosine triphosphatase (GTPase) activating protein (GAP) (SPAR), which has been “primed” for Plk2 binding by CDK5, a proline-directed kinase (Seeburg et al., 2008). The Plk2-SPAR binding results in proteasome-directed SPAR degradation, which has downstream effects on actin dynamics and Rap signaling, eventually leading to AMPAR and NMDAR removal and loss of spines. Whereas the Plk2-SPAR association is linked to synaptic AMPAR reduction without any reported preference to GluA1 or GluA2 subunits, a separate kinase-independent pathway in which Plk2 binds to N-ethylmaleimide-sensitive fusion protein (NSF) which is selective to GluA2 subunit removal has been observed (Evers et al., 2010).

Although there is evidence that excitatory synapses scale in a cell-wide, uniform manner during HSD, theoretical consideration invoke the existence of additional local dendritic mechanisms to assure proper information processing (Rabinowitch and Segev, 2006a,b). In fact, several local dendritic mechanisms which are engaged during homeostatic plasticity have been recently described. For example, chronic inactivity with NMDAR inhibition leads to retinoic acid signaling and stimulation of local GluA1 synthesis (Aoto et al., 2008; Poon and Chen 2008). Similarly, homeostatic upscaling via tetrodotoxin (TTX) and AMPAR/NMDAR blockade has also been shown to stimulate local protein synthesis (Sutton et al., 2007), including GluA1 accumulation (Sutton et al., 2006). In the context of HSD, miR-134-mediated local translation of Pumilio-2 (Pum2) mRNA upstream of Plk2 activation was reported (Fiore et al., 2014). The involvement of other miRNAs, namely miR-129 (Rajman et al., 2017) and miR-485 (Cohen et al., 2011) in HSD further suggest the importance of local translation in this process. Additionally, dendrite-specific regulation of excitatory proteins in the context of HSD has been demonstrated on a multi-omics scale (Colameo et al., 2021). Indeed, such local translational mechanisms could explain the spatial specificity of HSD, as demonstrated by homeostatic regulation in individual dendritic compartments (Rabinowitch and Segev, 2006a). The spatial specificity further extends to the synapse level, as scaling depends on the spatial patterns of synaptic potentiations (Rabinowitch and Segev, 2006b). Considering the mounting evidence for local regulation of proteins in plasticity mechanisms such as long-term potentiation (LTP) in both in vitro and in vivo contexts (Miller et al., 2002; Lyles et al., 2006), there is a need for identifying other proteins that are locally regulated in HSD.

In the present study, we investigate the expression regulation and function of Proline-rich 7/Transmembrane Adapter Protein 3 (Prr7) in the context of HSD induced by chronic activity. Prr7 is localized in neuronal dendrites of hippocampal neurons (Murata et al., 2005; Kravchick et al., 2016; Lee et al., 2018) and the postsynaptic density in rodent brains (Murata et al., 2005; Jordan et al., 2004; Yoshimura et al., 2004), suggesting that it could play an important role in HSD. Functionally, exosomally secreted Prr7 induces synapse elimination in hippocampal neurons (Lee et al. 2018), whereas NMDA-receptor mediated induction of excitotoxicity is accompanied by a translocation of Prr7 from the synapse to the nucleus, followed by a triggering of Jun-dependent apoptotic pathway (Kravchick et al., 2016). However, whether synaptically localized Prr7 is involved in activity-dependent forms of synaptic plasticity, e.g. HSD, is unknown.

Here, we report that the downregulation of Prr7 at both RNA and protein levels is required for dendritic spine elimination during HSD induced by chronic activity, and that the dendritic reduction of Prr7 is regulated post-transcriptionally by miR-329 and miR-495. Furthermore, our results suggest that the miR-329/495/Prr7 interaction ties in with the previously described SPAR/CDK5 pathway involved in homeostatic plasticity.

## Results

### Prr7 mRNA and protein are downregulated locally in processes by chronic activity

Prr7 has previously been identified as a synaptic protein and implicated in the control of excitatory synapse formation, but its role in synaptic plasticity, e.g. HSD is unknown. Specifically, whether the subcellular expression of Prr7 changes during HSD has not been investigated. To determine if Prr7 expression is regulated in HSD, we first examined Prr7 mRNA expression levels in mature (DIV21) primary rat hippocampal cells treated with either Mock (Ethanol) or the GABA-A receptor antagonist Picrotoxin (PTX) for 48h. This is a well established experimental paradigm to induce HSD in vitro, as demonstrated by expected changes in spike frequency, EPSC amplitude, and expression of the GluA1 subunit of AMPARs (Fiore et al., 2014, Rajman et al., 2017; Seeburg et al., 2008; Bateup et al., 2013; Evers et al., 2010; Ibata et al., 2008). Using qPCR, we observed a decrease in Prr7 mRNA levels in whole cell extracts of PTX-compared to Mock treated hippocampal neurons (Fig. 1a), which was similar in magnitude to GluA1 mRNA which was previously shown to be downregulated during HSD.

**Figure 1.**
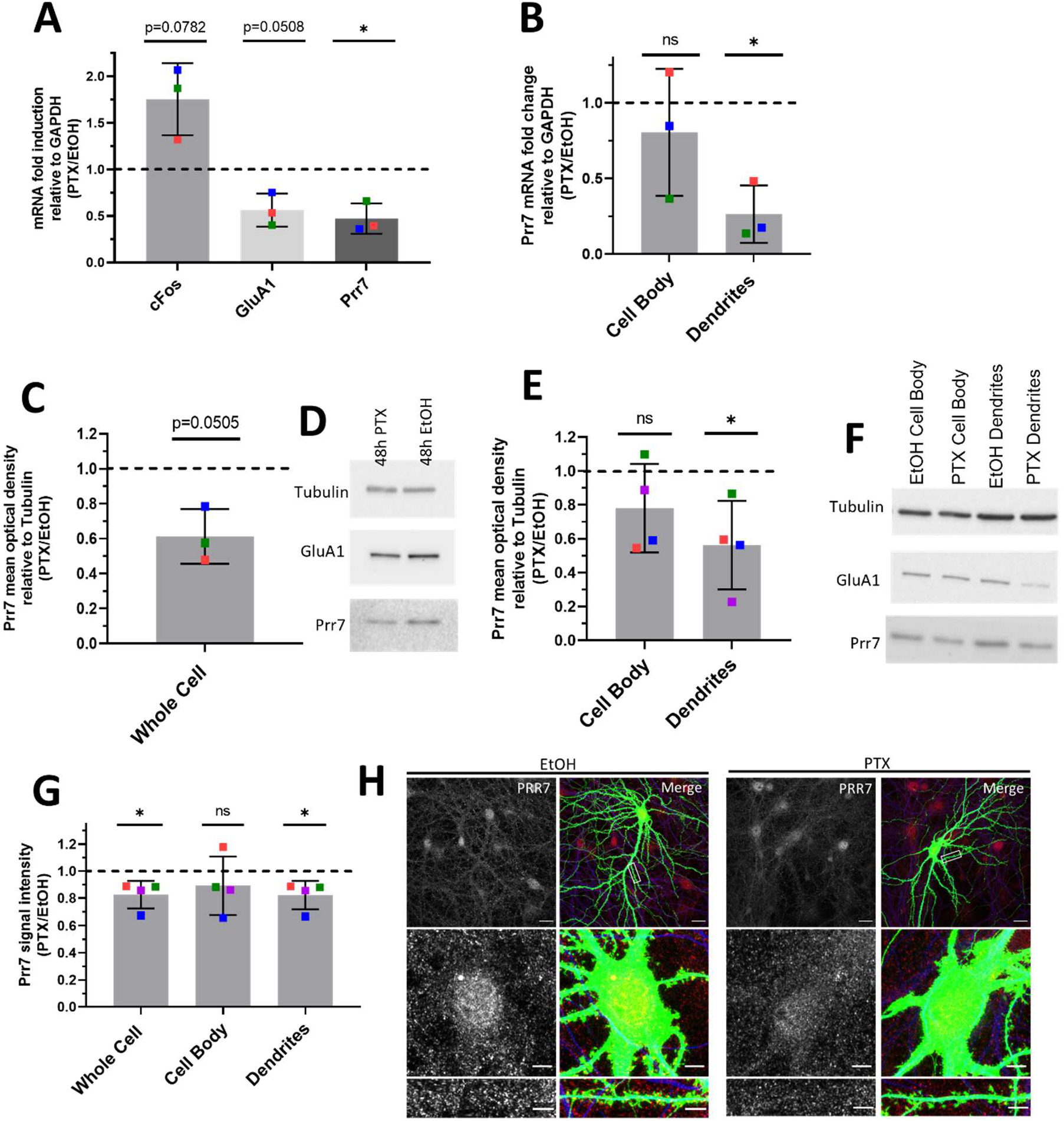
Global and local Prr7 downregulation at both RNA and protein levels by chronic activity. **(a)** Prr7 mRNA levels relative to GAPDH in hippocampal rat neurons treated with 100μM PTX or EtOH (1:500 volume) at DIV17 for 48h and lysed for RNA extraction on DIV19. * p=0.0303. **(b)** Prr7 mRNA levels in compartmentalized hippocampal cell samples treated at DIV19 with 100μM PTX or EtOH (1:500 volume) for 48h (ns p=0.5059, * p=0.0216). Prr7 protein levels as measured by mean optical band density relative to Tubulin in **(c-d)** whole cell hippocampal cell samples, and **(e-f)** compartmentalized hippocampal rat cultures treated at DIV19 with 100μM PTX or EtOH (1:500 volume) for 48h (ns p=0.1910, * p=0.0441). All replicate blots are shown in Supplemental Data. **(g)** Average Prr7 punctum intensity in GFP-transfected (150ng) cell body or dendrites selection of hippocampal rat neurons treated with PTX or EtOH on DIV19 for 48h. Each point represents the grand average for the 7-9 cells imaged in a single experiment (* p=0.0417 (whole cell), ns p=0.3988 (cell body), * p=0.0427 (dendrites)). **(h)** Representative whole cell, cell body, and dendrite images showing Prr7 expression (greyscale, left panels) and merged Prr7 (red), GFP (green), Map2 (blue) signals (right panels) in GFP-transfected hippocampal neurons treated with EtOH or PTX for 48h. Scale bars = 20μm (whole cell images) and 5μm (cell body and dendrite closeups). For all bar graphs, data = mean normalized to EtOH condition ±S.D., n=3-4, *p<0.05, one-sample t-test with hypothetical mean set to 1. Colors of points represent data from the same independent experiment.

Previous studies indicate that, in addition to neuron-wide changes, local alterations in gene expression in the synapto-dendritic compartment might also be involved in homeostatic plasticity (Colameo et al., 2021; Sutton et al., 2007). To determine local expression changes, we utilized a compartmentalized culture system as previously described (Bicker et al., 2013), which allowed separate measurements of Prr7 RNA expression in Mock vs. PTX-treated cells in cell bodies and processes (which are mainly represented by dendrites), respectively. Thereby we observed a significant decrease in Prr7 mRNA levels in the process compartment upon PTX treatment (Fig. 1b). Prr7 levels in the cell body compartment were more variable, but also trended downward by PTX, consistent with our observations in whole cell extracts.

We further probed for Prr7 protein expression by immunoblotting whole cell and compartmentalized protein extracts from PTX and Mock-treated hippocampal neurons. Whereas there was a general downward trend in Prr7 protein levels upon PTX treatment, the effect was most pronounced and statistically significant in dendrites, providing further support for an important contribution of local regulatory mechanism engaged in the control of Prr7 expression during HSD (Fig. 1c-f).

To further corroborate the observed subcellular differences in Prr7 regulation in neurons, we additionally analyzed Prr7 protein levels through Prr7 immunostaining of GFP-transfected hippocampal neurons which were either Mock- or PTX-treated. Therefore, we used a commercial Prr7 antibody whose specificity was validated by the presence of reduced signal intensity in Prr7 knockdown cells (Supplemental Fig. 1). We measured the average Prr7 puncta intensities within whole cell, cell body, and dendrite (whole cell with cell body removed) selections using GFP as a mask. Consistent with the Western blot data, reduction in Prr7 puncta intensity upon PTX was most robustly observed in neuronal dendrites, whereas analysis of cell bodies only revealed a non-significant reduction of the Prr7 signal (Fig. 1g-h). This decrease was homogenous along the dendrites since no difference in Prr7 downregulation was detected between proximal vs. distal dendrites (Supplemental Fig. 2).

Notably, no significant differences in neither Prr7 mRNA nor protein levels between the cell body and dendritic compartments were observed in Mock-treated neurons at baseline (Supplemental Fig. 3), indicating that mechanisms leading to Prr7 downregulation are specifically engaged during HSD.

### Downregulation of Prr7 is necessary and sufficient for spine density reduction during HSD

Based on our finding that Prr7 is downregulated during HSD, we asked whether Prr7 knockdown is sufficient to induce HSD. Prr7 knockdown was achieved using transfection of a Prr7 shRNA expressing plasmid based on a previously published Prr7 targeting sequence (Kravchick et al., 2016) We confirmed efficient and specific knockdown of Prr7 using the generated construct (Supplemental Fig. 4). Prr7 knockdown did not adversely affect cell health based on unaltered cell morphology between control and Prr7 shRNA-transfected neurons (Supplemental Fig. 1b).

Using the validated shRNA construct, we found that Prr7 loss in hippocampal neurons led to a significant decrease in dendritic spine density, to levels comparable to those induced by PTX (Fig. 2a-b). To determine whether spine density reduction was specific to Prr7 knockdown and not caused by off-target effects of the shRNA used, we generated an shRNA-resistant Prr7 expression construct (validations shown in Supplemental Fig 5 and 6). The spine density reduction from Prr7 shRNA was rescued when the shRNA-resistant Prr7 expression construct was introduced (Fig. 2c-d). Moreover, a small (~34%) increase in spine density was observed upon Prr7 overexpression alone. Furthermore, overexpression of Prr7 in hippocampal neurons led to complete prevention of spine density reduction in the presence of PTX (Fig. 2e-f).

**Figure 2.**
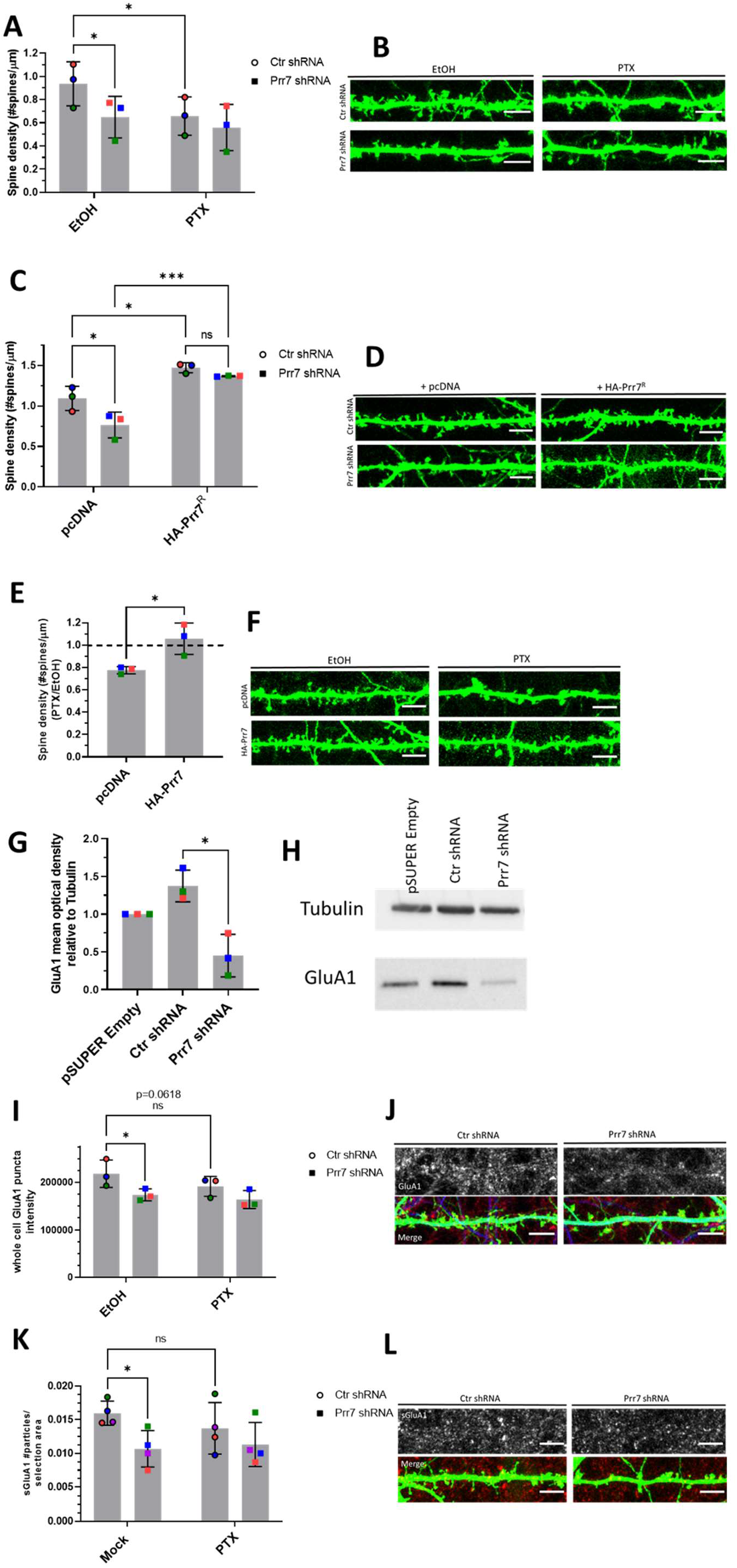
Prr7 downregulation is necessary and sufficient for HSD. **(a-b)** Spine densities of hippocampal rat neurons transfected with GFP (150ng) and either control or Prr7 shRNA vector (7.5ng pSUPER) on DIV13, treated at DIV19 with 100μM PTX or EtOH (1:500 volume) for 48h, with representative GFP images showing dendrites for each condition (Ctr EtOH vs. Prr7 EtOH: p=0.0175; Ctr EtOH vs. Ctr PTX: p=0.0187). **(c-d)** Spine densities of hippocampal neurons transfected with GFP (150ng), either control or Prr7 shRNA (7.5ng) and either pcDNA or shRNA-resistant Prr7 construct (400ng) on DIV13 and fixed on DIV20-21 (Ctr pcDNA vs. Prr7 pcDNA: p= 0.0309; Ctr pcDNA vs. Ctr HA-Prr7^R^: p=0.0146; Prr7 pcDNA vs. Prr7 HA-Prr7R: p=0.0008; Ctr HA-Prr7^R^ vs. Prr7 HA-Prr7^R^: p=0.6899). For these shRNA experiments, data = mean ±S.D., n=3, *p<0.05, *** p<0.001, two-way ANOVA with Tukey’s post-hoc HSD test. **(e-f)** Spine densities of hippocampal neurons transfected with GFP (150ng) and either pcDNA or HA-Prr7 construct (400ng) on DIV13 and treated at DIV19 with EtOH or PTX for 48h. Data = mean normalized to EtOH condition ± S.D., n = 3, * p=0.0280, unpaired Student’s t-test. **(g-h)** GluA1 protein levels as measured by mean optical band density relative to Tubulin in empty pSUPER, Ctr shRNA, and Prr7 shRNA-transfected cortical neuron whole cell extracts, with representative western blot images. Data = mean normalized to pSUPER empty condition ± S.D., n = 3, * p=0.0105, unpaired Student’s t-test. Full dataset of blots are shown in Supplemental Fig. 4. **(i-j)** Average whole cell GluA1 puncta intensity and **(k-l)** surface GluA1 puncta number for hippocampal neurons transfected with GFP, with either control or Prr7 shRNA, and treated with PTX or EtOH for 48h. Data = mean ± S.D, n=3, 2-way ANOVA with Tukey’s post-hoc HSD test. Whole cell GluA1: Ctr EtOH vs. Prr7 EtOH: * p=0.0228; Ctr EtOH vs. Ctr PTX: ns p=0.0618. Surface GluA1: Ctr EtOH vs. Prr7 EtOH: * p=0.0121; Ctr EtOH vs. Ctr PTX: ns p=0.1193. All scale bars shown = 5μm. Whole cell images from which dendrite segments were taken are shown in Supplemental Data.

Since AMPAR degradation is a hallmark of HSD, we studied the effect of Prr7 knockdown on the protein levels of the GluA1 subunit of AMPARs. Knockdown of Prr7 in cortical neurons led to a reduction in total GluA1 protein levels as judged by immunoblotting (Fig. 2g-h). Additionally in hippocampal neurons, GluA1 whole cell puncta intensity and surface GluA1 puncta number were reduced in the Prr7 knockdown condition based on immunostaining (Fig. 2i-l). Together, these observations suggest that loss of Prr7 might be sufficient to induce a reduction in spine density and AMPA-type glutamate receptor subunits, both of which are commonly observed during HSD.

### miR-329 and miR-495 are required for Prr7 downregulation by PTX

Next, we explored the mechanisms underlying PTX-dependent downregulation of Prr7. Following the observations that Prr7 expression is regulated at the RNA level upon PTX treatment, we hypothesized that it may be regulated post-transcriptionally by miRNAs. miRNAs already have strong implications toward activity-dependent synaptic plasticity mechanisms, including HSD (Fiore et al., 2014; Cohen et al., 2011; Rajman et al., 2017).

Upon analysis of the Prr7 3’ UTR sequence using the Targetscan algorithm, we found four predicted miRNA binding sites, two of which overlap with one another (Fig. 3a). We examined the effect of inhibiting two of the four miRNA candidates, miR-329-3p and miR-495-3p, on Prr7 mRNA expression in the context of PTX treatment, through the use of a luciferase reporter with the Prr7 3’ UTR cloned downstream of a firefly gene. We selected these two miRNAs for further studies since miR-495-3p is the most abundant of the four candidates, and miR-329-3p has been previously implicated in KCl-dependent dendritogenesis (Fiore et al., 2009). We observed a reduction in firefly luciferase activity upon PTX stimulation, which was prevented by a cocktail of miR-329-3p and miR-495-3p inhibitors (antisense locked nucleic acid inhibitors “pLNAs”). Importantly, this effect was not seen when a reporter with mutated binding sites for these miRNAs on the Prr7 3’ UTR was used (Fig. 3b), demonstrating that the effects were mediated by the miRNA binding sites present in the Prr7 3’UTR. The same effect was observed when transfecting miR-495-3p pLNA alone (Fig. 3c). A trend was observed for miR-329-3p pLNA alone, although the effect did not reach statistical significance (Fig. 3d). Taken together, these findings indicated that Prr7 is a direct target of miR-329-3p and miR-495-3p, with miR-495-3p contributing more to the downregulation, and that these interactions take place during HSD.

**Figure 3.**
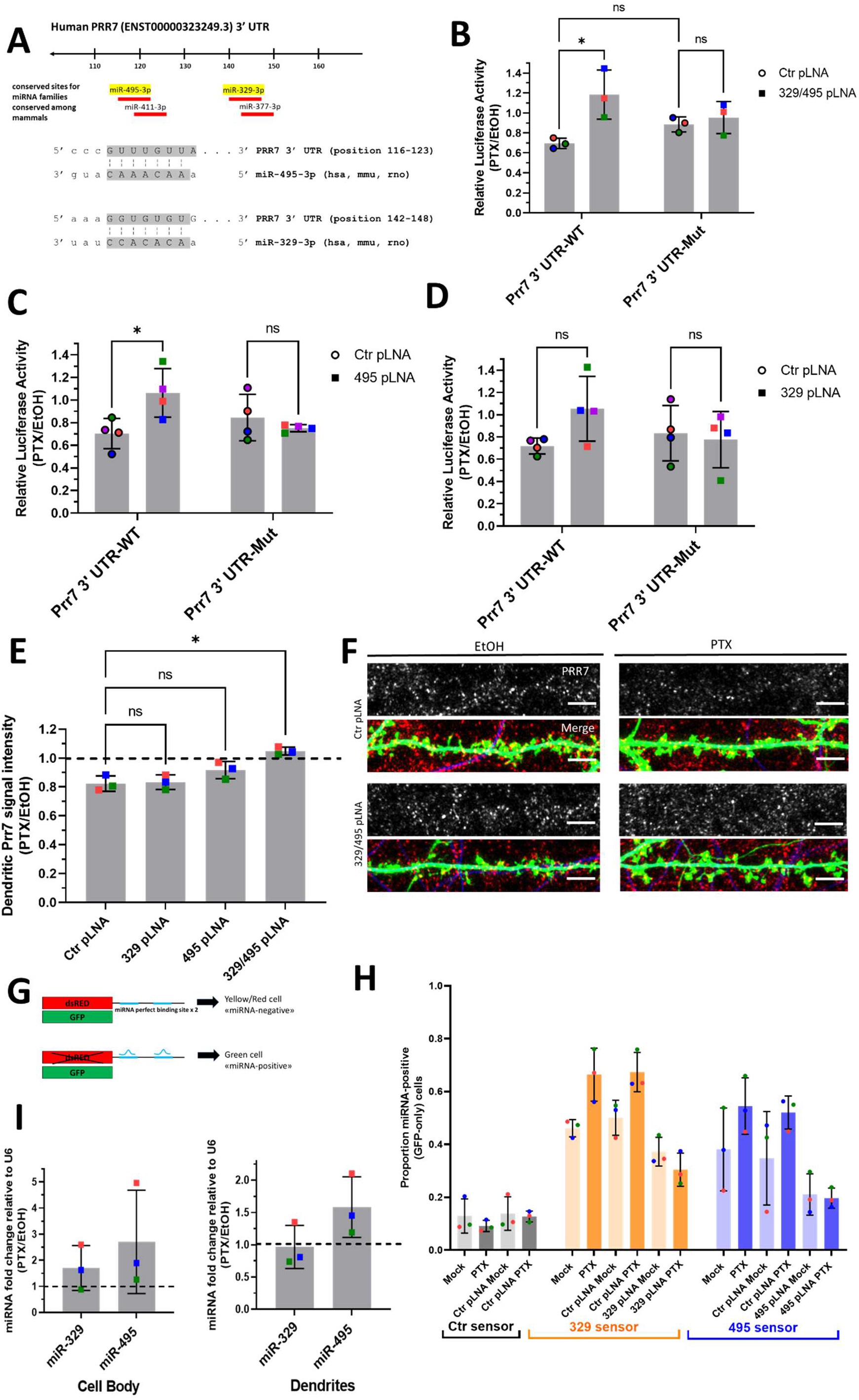
miR-329 and miR-495 are required for Prr7 downregulation by PTX. **(a)** Predicted miRNA binding sites for human Prr7 3’ UTR, and seed matches for miR-329 and miR-495 from Target Scan (http://www.targetscan.org/). **(b-d)** Rat hippocampal neurons were co-transfected at DIV13 with either pmiRGLO Prr7 WT or Mut 3’ UTR plasmids (50ng), and control pLNA, 329 pLNA, 495 pLNA (20pmol) or miR-329/495 pLNA mix (10pmol each), and treated with either EtOH or 100μM PTX at DIV18. Cells were lysed at DIV20 and firefly/renilla luciferase activity ratios measured. Data = mean normalized to EtOH condition ± S.D, n=3-4, 2-way ANOVA with Tukey’s post-hoc HSD test. 329/495 pLNA data: Ctr WT vs. 329/495 WT: * p=0.0194; Ctr WT vs. Ctr Mut: ns p=0.4724; Ctr Mut vs. 329/495 Mut: ns p=0.9452. 495 pLNA data: Ctr WT vs. 495 WT: * p=0.0397; Ctr Mut vs. 495 Mut: ns p=0.8484. 329 pLNA data: Ctr WT vs. 329 WT: ns p=0.2275; Ctr Mut vs. 329 Mut: ns p=0.9841. **(e)** Average Prr7 punctum intensity in hippocampal cells transfected with GFP (150ng) and control, miR-329, miR-495 pLNA (20pmol), or miR-329/495 pLNA mix (10pmol each) at DIV13 and treated with PTX or EtOH on DIV19 for 48h. Data = mean normalized to EtOH condition ± S.D., n = 3, one-way ANOVA with Tukey’s post-hoc HSD test. Ctr vs. 329: ns p=0.9946; Ctr vs. 495: ns p=0.1739; Ctr vs. 329/495: * p=0.0023. **(f)** Representative dendrite images showing Prr7 expression (grey-scale, top panels) and merged Prr7 (red), GFP (green), Map2 (blue) signals (bottom panels) in Ctr or 329/495 pLNA-transfected hippocampal neurons treated with 48h EtOH or PTX. Scale bars = 5μm. Whole cell images from which dendrite segments were taken are shown in Supplemental Data. **(g-h)** Schematic of single-cell dual-fluorescence miRNA sensor assay, and measurement of endogenous miR-329 and miR-495 activity upon PTX treatment in hippocampal neurons. Cells were transfected with miR-329, miR-495, or control sensor (125ng), with or without Ctr, miR-329, or miR-495 pLNA (5pmol) at DIV13, treated with 100μM PTX or EtOH at DIV19, and fixed at DIV21. The number of neurons expressing GFP only without dsRed vs. those expressing dsRed were counted. Data = mean proportion of GFP+ cells/total cell count ± S.D, n=3. **(i)** Mature miR-329 and miR-495 levels in cell body and dendrite compartments of hippocampal neurons treated with either EtOH or PTX for 48h. Data = mean normalized to EtOH condition ± S.D., n = 3.

We then asked whether the miR-329-3p and miR-495-3p regulation of Prr7 during downscaling as suggested by luciferase could also be seen at the protein level for dendrite-localized Prr7. Decreases in Prr7 in dendrites upon PTX were prevented when cells were transfected with the cocktail of miR-329-3p and miR-495-3p pLNAs (Fig. 3e-f), but not for miR-329-3p or miR-495-3p pLNA alone, thereby confirming our results from luciferase assays and indicating an additive inhibitory role for miR-329-3p and miR-495-3p in Prr7 regulation.

Since miRNA inhibition appeared to upregulate Prr7 only in the context of PTX stimulation, we speculated that miR-329-3p and miR-495-3p themselves could be subject to PTX-dependent regulation. We examined endogenous miRNA activity in the Mock and PTX-stimulated hippocampal neurons through use of a single cell dual fluorescence assay (“sensor assay”), as previously described (Fiore et al., 2009). Specifically, the assay utilizes polycistronic vectors expressing both GFP and dsRed, whereby dsRed expression is post-transcriptionally controlled by the presence of two perfectly complementary binding sites for the miRNA of interest within the dsRed 3’ UTR (Fig. 3g). If miRNAs of interest are active within a given cell, they would bind to the dsRed 3’UTR and downregulate dsRed expression. Thus, cells expressing only GFP without dsRed were counted as “miRNA positive”, and those expressing dsRed were counted as “miRNA negative”. Hippocampal neurons were transfected either with a control sensor (containing a sequence non-specific to any known miRNAs), a miR-329-3p or a miR-495-3p sensor. Subsequently, “miRNA positive” vs. “miRNA negative” cells were manually scored over the entirety of each coverslip for all conditions (Supplemental Fig. 7). The proportion of miRNA-positive neurons increased upon PTX treatment for both the miR-329-3p and miR-495-3p sensor transfections. This induction was not seen when a pLNA against the respective miRNA was co-transfected with the sensor, indicating that the sensor could reliably detect endogenous miRNA activity. In conclusion, PTX treatment increased the proportion of neurons displaying active miR-329-3p and miR-495-3p (Fig. 3h).

Next, we wanted to test whether the observed PTX-dependent increase in miR-329/495 activity was due to an upregulation of miRNA expression. qPCR analysis of these two miRNAs in RNA extracts obtained from compartmentalized neuron cultures indicated a non-significant PTX-dependent upregulation in mature miRNA levels for both miR-329-3p and miR-495-3p in the cell body upon PTX treatment (Fig. 3i). In the process compartment, mature miR-495-3p, but not miR-329-3p levels were significantly increased by PTX. These findings suggest that in the case of miR-495-3p, PTX-dependent activity increase might involve a local upregulation of miR-495-3p expression in the dendritic compartment.

### miR-329-3p and miR-495-3p are required for synaptic depression induced by PTX and Prr7 knockdown

We next asked whether miR-329-3p and miR-495-3p were functionally involved in HSD. Therefore, we measured spine density in cells transfected with miR-329-3p and miR-495-3p pLNAs in the presence or absence of PTX treatment (48h). We found that both miR-329-3p and miR-495-3p inhibition, separately and together, rescued PTX-mediated spine density reduction (Fig. 4a-b). The rescue effect was most pronounced when using a miR-329/495 pLNA cocktail, consistent with our results from Prr7 regulation (Fig. 3b, e).

**Figure 4.**
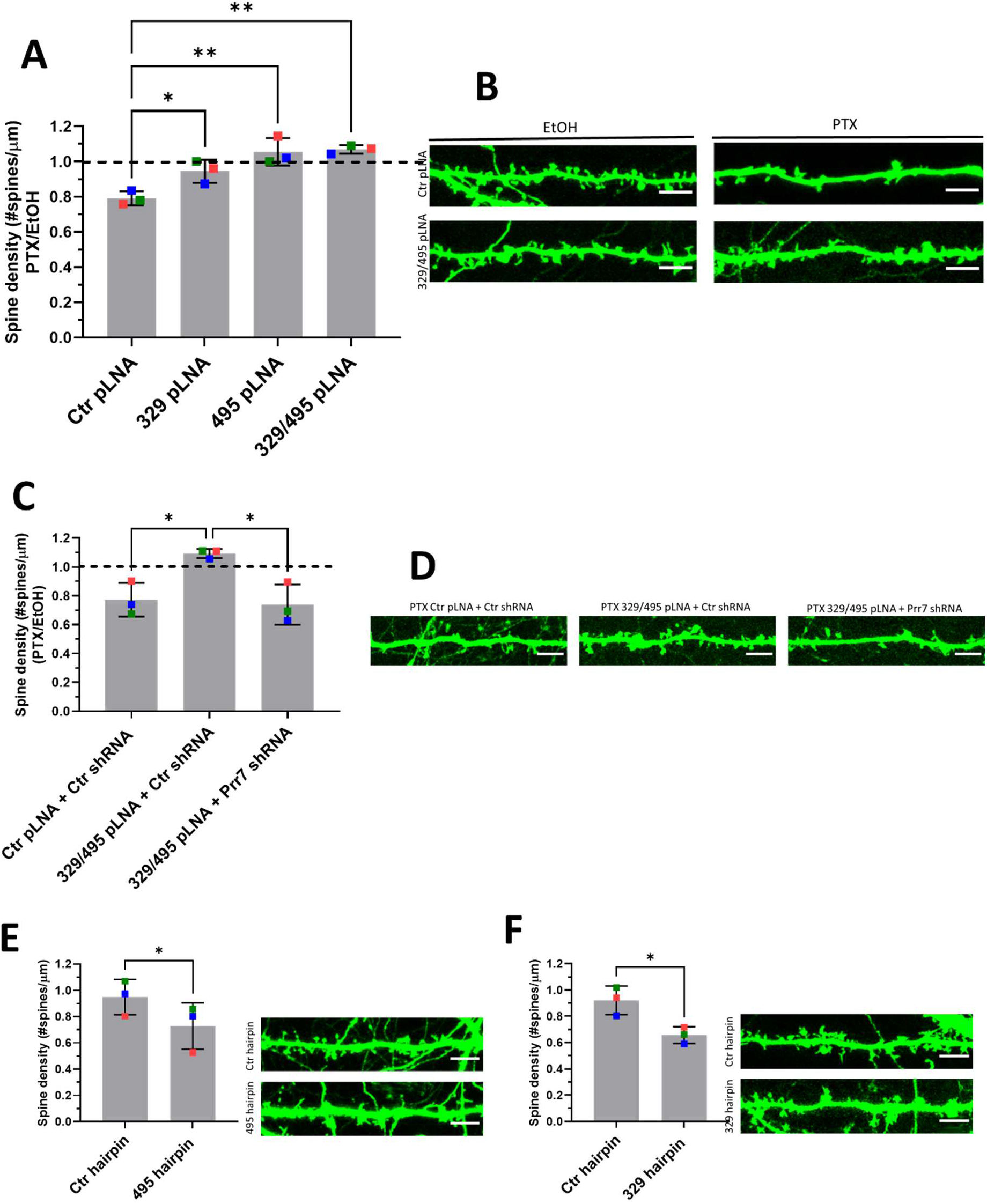
miR-329 and miR-495-mediated Prr7 downregulation are required for synaptic depression induced by PTX. **(a-b)** Spine densities of hippocampal cells transfected with GFP (150ng) and control, miR-329, miR-495 pLNA (20pmol) or miR-329/495 pLNA mix (10pmol each) on DIV13, treated with EtOH or 100μM PTX on DIV19 for 48h. Representative GFP images showing dendrites for each condition are shown. Ctr vs. 329: * p=0.0400; Ctr vs. 495: ** p=0.0018; Ctr vs. 329/495: ** p=0.0013. **(c-d)** Spine densities of hippocampal cells transfected with GFP (150ng), control pLNA (20pmol) or miR-329/495 pLNA mix (10pmol each), and control or Prr7 shRNA (2.5ng pSUPER) on DIV13, treated with EtOH or PTX on DIV19 for 48h (Ctr + Ctrsh vs. 329/495 + Ctrsh: * p=0.0240; 329/495 + Ctrsh vs. 329/495 + Prr7sh: * p=0.0155). For these pLNA data, data = mean normalized to EtOH condition ±S.D., n=3, one-way ANOVA with Tukey’s post-hoc HSD test. Spine densities of hippocampal cells transfected with control or **(e)** miR30a-495 chimeric hairpin (500ng), **(f)** miR30a-329 chimeric hairpin (500ng) on DIV13 and fixed on DIV18-19 (329hp) or DIV21 (495hp). Data = mean ± S.D., n=3, paired Student’s t-test (Ctr vs. 495: * p=0.0195; Ctr vs. 329: * p=0.0285). Representative GFP images showing dendrites for each condition are shown. Whole cell images from which dendrite segments were taken are shown in Supplemental Data.

To corroborate Prr7 as an important downstream target in miR329/495-mediated HSD, we further asked whether the impaired HSD induced by the pLNA cocktail could be reinstated by lowering Prr7 levels through co-transfection of Prr7 shRNA. Consistent with this idea, transfection of Prr7 shRNA, but not control shRNA, restored the PTX-induced spine density reduction in the presence of miR-329 and miR-495 pLNAs (Fig. 4c-d). This result demonstrates that Prr7 is a key target of miR-329/-495 in PTX-mediated HSD.

We went on to test whether increasing levels of miR-329 and −495 was sufficient to induce spine elimination in the absence of PTX, thereby mimicking HSD. Towards this end, we constructed chimeric miR-329 and −495 overexpressing plasmids using a previously described strategy (Christensen et al. 2010; hairpin diagram Supplemental Fig. 8) which allows for efficient miRNA overexpression. Roughly 2-fold overexpression of the miRNA of interest relative to control (Supplemental Fig. 8) was observed, accompanied by consistently downward trends of Prr7 in dendrites (Supplemental Fig. 9). Despite the moderate effects on miRNA overexpression and Prr7 reduction achieved with this approach, stable overexpression of miR-329 and miR-495 was in both cases sufficient to induce a significant reduction in spine density (Fig. 4e-f). Thus, miR-329/495 overexpression mimics PTX-induced miR-329/495 expression followed by spine elimination in transfected hippocampal neurons.

### SPAR/CDK5 pathway is downstream of miR-329/miR-495/Prr7 in HSD

We further explored the pathway downstream of miR329/495/Prr7 which mediates the effects on spine density. One attractive candidate is the Plk2/SPAR pathway which has previously been implicated in HSD (Pak and Sheng, 2003). Specifically, Plk2-mediated phosphorylation of SPAR is followed by proteasome-dependent SPAR degradation, leading to excitatory synapse weakening and spine loss. Intriguingly, both SPAR and Prr7 have been shown to interact with PSD-95, an important scaffold protein required for the integrity of the post-synaptic density. We therefore speculated that Prr7 might protect SPAR from Plk2-mediated degradation, possibly in conjunction with PSD-95. Thus, we tested whether reducing Prr7 levels affected SPAR expression in a way consistent with a role in HSD. We found a reduction in SPAR levels in cortical neurons nucleofected with Prr7 shRNA through western blotting (Fig. 5a-b). Moreover, dendrite-localized SPAR protein was reduced in Prr7 shRNA-transfected hippocampal neurons (Fig. 5c-d; Supplemental Fig. 11). SPAR reduction was also seen upon PTX treatment as previously reported (Pak and Sheng, 2003). Therefore, Prr7 may stabilize SPAR at basal levels of network activity.

**Figure 5.**
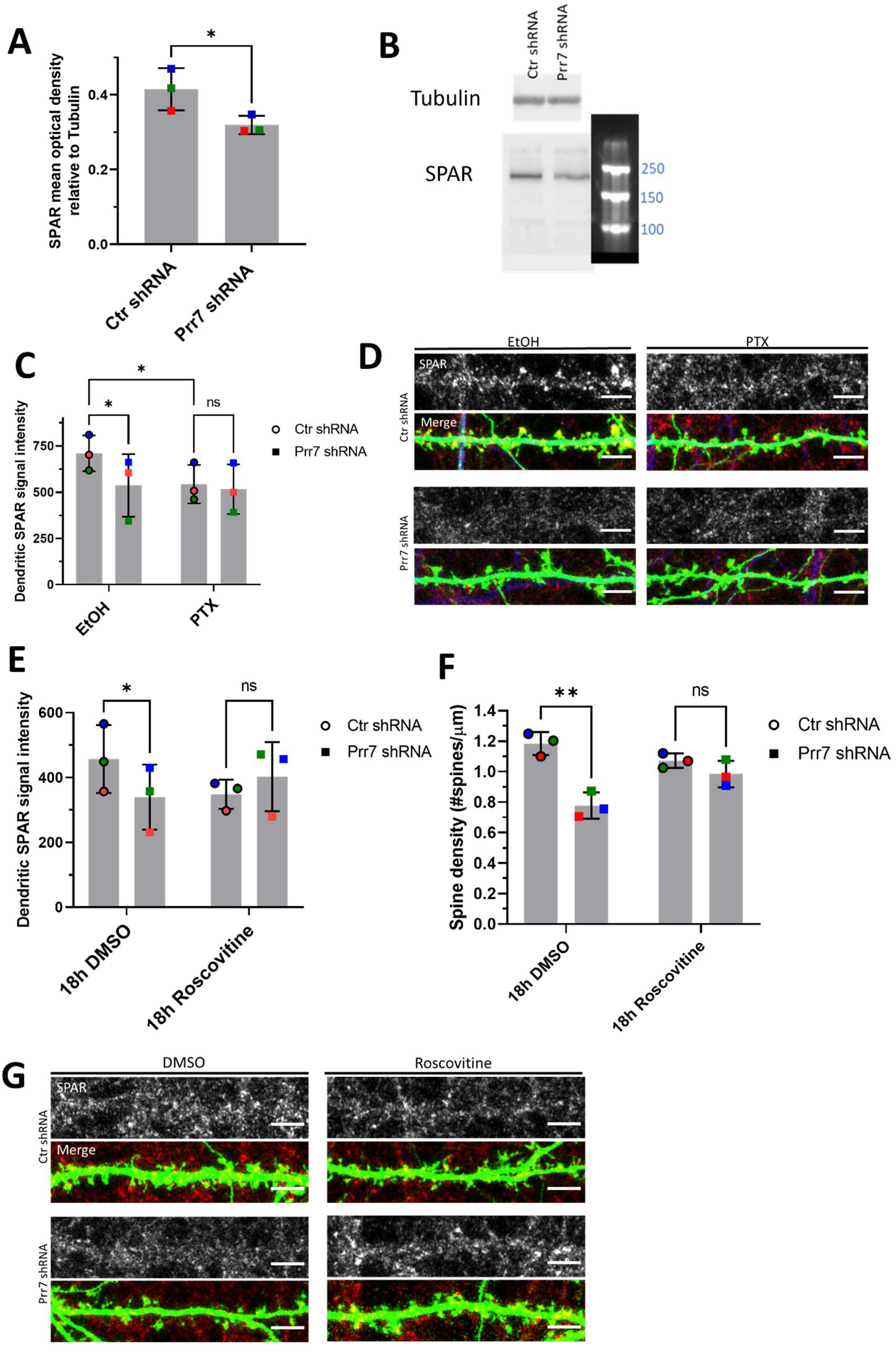
SPAR/CDK5 pathway is downstream of miR-329/495/Prr7 regulation in HSD. **(a-b)** SPAR protein levels relative to Tubulin in rat cortical neurons nucleofected with control or Prr7 shRNA (2μg) on day of dissociation (E18) and harvested for protein extraction 5 days later, with representative western blot (all replicate blots are shown in Supplemental Data). Data = mean ± S.D., n=3, * p=0.0462, paired Student’s t-test. **(c)** Average SPAR punctum intensity in dendrite selection in rat hippocampal cells transfected with GFP (150ng), control or Prr7 shRNA (7.5ng pSUPER) at DIV13, treated with either EtOH (1:500 volume) or 100μM PTX at DIV19 for 48h, then immunostained for SPAR. Data = mean ± S.D., where each point represents grand average in dendrites for the 7-10 cells imaged in a single experiment. n=3, 2-way ANOVA with Tukey’s HSD post-hoc test. Ctr EtOH vs. Prr7 EtOH: * p= 0.0429; Ctr EtOH vs. Ctr PTX: * p=0.0462; Ctr PTX vs. Prr7 PTX: ns p=0.6932. **(d)** Representative dendrite images showing SPAR expression (grey-scale, top panels) and merged SPAR (red), GFP (green), Map2 (blue) signals (bottom panels) in Ctr or Prr7 shRNA-transfected hippocampal neurons treated with 48h EtOH or PTX. **(e)** Average dendrite SPAR punctum intensity in rat hippocampal cells transfected with GFP (150ng) and either control or Prr7 shRNA (7.5ng) at DIV13, treated with either DMSO (1:1000 volume) or Roscovitine (10μM) at DIV19 for 18h, then immunostained for SPAR. Data = mean ± S.D., where each point represents grand average in dendrites for the 8-12 cells imaged in a single experiment. n=3. Ctr DMSO vs. Prr7 DMSO: * p=0.0266; Ctr Ros vs. Prr7 Ros: ns p=0.2328. **(f)** Spine densities for these same cells were measured. Ctr DMSO vs. Prr7 DMSO: ** p=0.0064; Ctr Ros vs. Prr7 Ros: ns p=0.4288. For these Roscovitine data, data = mean ± S.D., 2-way ANOVA with Tukey’s HSD post-hoc test. **(g)** Representative dendrite images showing SPAR expression (grey-scale, top panels) and merged SPAR (red) and GFP (green) signals (bottom panels) in Ctr or Prr7 shRNA-transfected hippocampal neurons treated with 18h DMSO or Roscovitine. Scale bars = 5μm. Whole cell images from which dendrite segments were taken are shown in Supplemental Data.

CDK5 kinase-mediated SPAR phosphorylation primes SPAR for targeting by Plk2 (Seeburg et al., 2008). To determine whether Prr7 serves to stabilize SPAR by interfering with CDK5 activity, we treated Prr7 shRNA-transfected cells with 10μM Roscovitine, a CDK5 inhibitor, and quantified SPAR puncta intensity in dendrites. We found that a reduction in SPAR protein levels was no longer seen in Prr7 shRNA-transfected cells treated with Roscovitine relative to DMSO (Fig. 5e, g). Moreover, Roscovitine treatment rescued the spine density reduction in the Prr7 knockdown condition (Fig. 5f-g). These findings suggest that Prr7 functions upstream of CDK5, potentially stabilizing dendritic spines through protecting SPAR from CDK5-mediated priming phosphorylation, which in turn is required for SPAR phosphorylation by Plk2 and subsequent proteasome-dependent degradation.

## Discussion

### Summary

Our study demonstrates the requirement of miR-329 and miR-495-mediated downregulation of Prr7 underlying dendritic spine elimination in HSD. From our results, we present the following model (Fig. 6). Under basal conditions, Prr7 mRNA is actively translated, as the targeting of the Prr7 3’ UTR by miRNAs is inhibited by a yet unknown mechanism. Prr7 protein is required for the stabilization of SPAR through inhibiting the activity of CDK5, thereby maintaining the integrity of the post-synaptic density, including the stabilization of GluA1-containing AMPARs at the surface.

**Figure 6.**
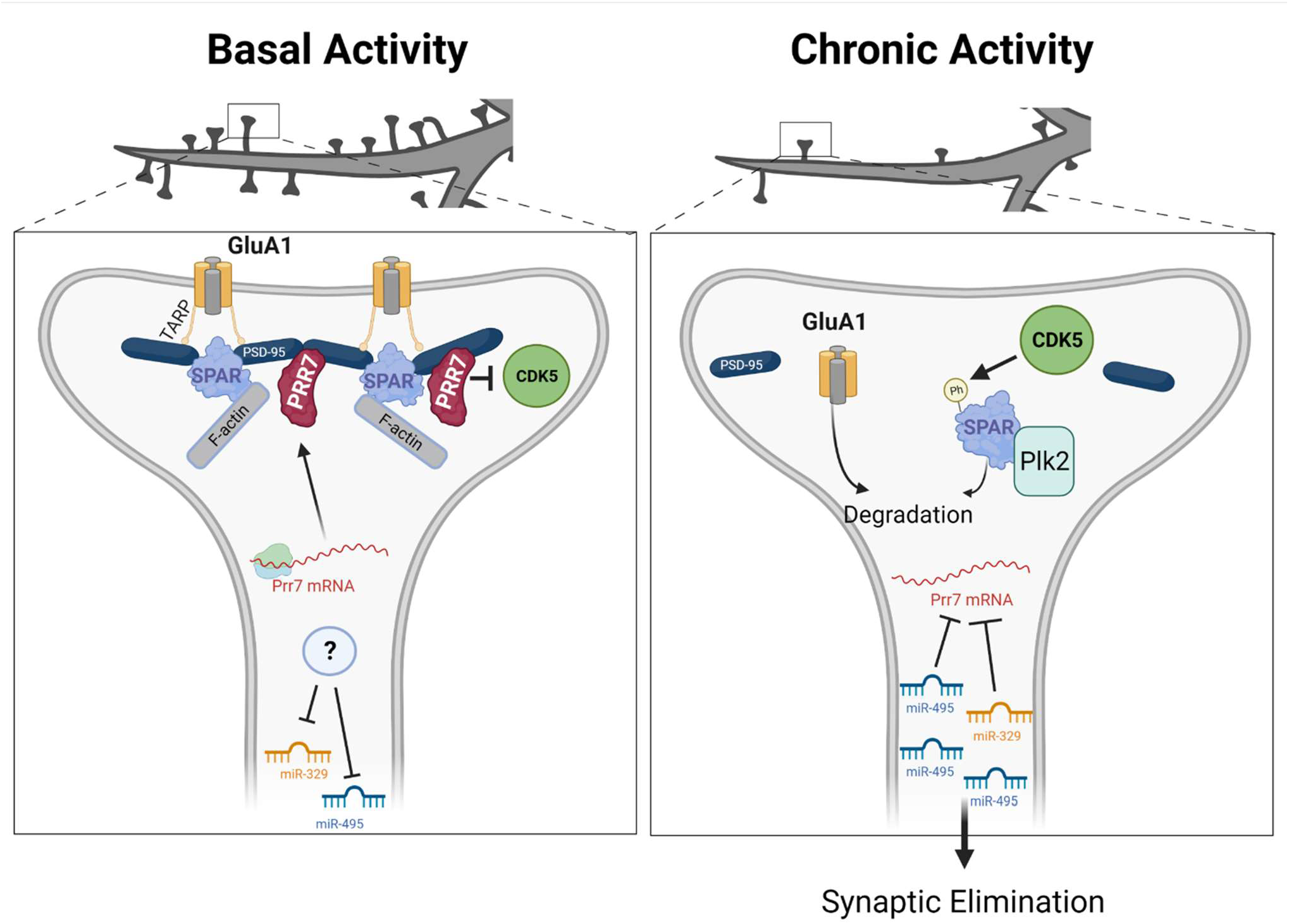
Model of miRNA-mediated Prr7 downregulation and downstream effects on SPAR during HSD. Left panel: under basal conditions, Prr7 mRNA is actively translated, as the miR-329 and miR-495 activity are inhibited (by a yet unknown mechanism). Prr7 protein stabilizes SPAR through inhibiting CDK5 activity and preventing SPAR phosphorylation. Thereby, the integrity of the post-synaptic density is maintained. Right panel: following chronic activity, miR-329 and miR-495 are activated, and miR-495 expression specifically is increased in dendrites. These miRNAs inhibit Prr7 mRNA, and thus Prr7 protein is lost. CDK5 phosphorylates SPAR, leading to targeting of SPAR by Plk2, and subsequent proteasomal degradation. As a result of SPAR loss, PSD-95 complexes are destabilized and GluA1 is degraded, leading to elimination of mature spines. (Figure generated with BioRender).

In contrast, following chronic activity, miR-329 and miR-495 are activated, and miR-495 expression is increased in dendrites. These miRNAs repress translation of Prr7 mRNA. In the absence of Prr7 protein, CDK5 phosphorylates SPAR, leading to an association between SPAR and Plk2, and subsequent proteasomal SPAR degradation. The loss of SPAR results in the destabilization of PSD-95 complexes and GluA1 degradation, ultimately resulting in spine elimination.

### Role of Prr7 in synaptogenesis and plasticity

Through Prr7 knockdown studies, we have revealed that Prr7 reduction leads to a decrease in spine number (Fig. 2a) as well as GluA1 protein levels (Fig. 2h-k), recapitulating two hallmarks of HSD. Although Prr7 was not found to associate with AMPARs in a previous study through immunoprecipitations (Kravchick et al., 2016), it is still possible that Prr7 influences AMPAR dynamics indirectly through interaction with other PSD components, e.g. the AMPAR auxiliary subunit Stargazin, which binds to both AMPARs and PSD-95 (Bats et al., 2007). Nevertheless, the current results are in agreement with the previously presented idea that Prr7 reduction serves a neuroprotective function against over-excitation (Kravchick et al., 2016), as well as Prr7 forms part of the post synaptic density core to promote neuronal maturation (Murata et al., 2005). The direct and indirect protein interactions involving Prr7, and Prr7-associated complexes formed under basal vs. stimulated conditions need further clarification.

Our results using sparse transfection of hippocampal neuron cultures clearly indicate a cell-autonomous, postsynaptic function of Prr7. In contrast, a non-cell autonomous function of Prr7 through exosomal secretion and Wnt inhibition has previously been reported (Lee et al., 2018). In this study, treatment of exosomal Prr7-rich supernatant in hippocampal cultures led to a reduction of glutamatergic synapses, which is contrary to our observations. The differences in incubation time between treatment and imaging (fixation 18h vs. 8 days post-transfection) may account for this discrepancy. Namely, it is possible that upon acute (18h) Prr7 overexpression, spines are eliminated due to a rapid exosomal Prr7 secretion from the soma. In contrast, over a time scale of days, Prr7 might accumulate in the synapto-dendritic compartment where it promotes synaptogenesis to compensate for the initial spine loss. In addition, the observed increase in excitatory synapses upon Prr7 knockdown in Lee et al., was solely based on PSD-95 puncta number, and effects on spines were not addressed.

We further elucidated a mechanism in which CDK5/SPAR is controlled downstream of Prr7 activity (Fig. 5a-g). It is interesting to consider this new pathway linking miR-329/495/Prr7 to SPAR in relation to a study describing miR-134-dependent SPAR regulation via Pum2 downregulation and Plk2 function (Fiore et al., 2014). It is known that upon chronic activity Plk2 is activated, and there is a bifurcation into two downstream branches (Seeburg and Sheng, 2008; Seeburg et al., 2008; Evers et al., 2010): 1) activated Plk2 phosphorylates SPAR, leading to GluA1/GluA2 internalization, and 2) activated Plk2 phosphorylates the GluA2-interacting protein NSF, promoting specifically GluA2 internalization. Intriguingly, in Fiore et al. it was found that the miR-134 pathway only affected GluA2 levels, and therefore it was suggested to connect only to the second branch. Considering Prr7 knockdown affected GluA1 expression (Fig. 2h-i), it would be plausible that conversely miR-329/495/Prr7 feeds into specifically the first branch via influencing CDK5.

### Role of miRNAs in local activity-dependent regulation of Prr7 during HSD

We have demonstrated that Prr7 expression at both RNA and protein levels is consistently reduced in the dendritic compartment in response to chronic activity (Fig 1b, e, g). The decrease in Prr7 mRNA in dendrites of PTX-treated neurons was also observed from RNA-seq analyses (Colameo et al., 2021). These observations support the idea that there is local regulation of Prr7 in the synapto-dendritic compartment during HSD. In this regard, local homeostatic mechanisms at the level of individual dendritic domains (Ju et al., 2004; Sutton et al., 2006) and at individual synapses (Hou et al., 2008) have been previously demonstrated. The exact mechanism by which local downregulation of Prr7 occurs is yet to be uncovered and will need the employment of techniques that allow the visualization of newly synthesized proteins. These include for example puromycin labeling with proximity ligation assay (puro-PLA) (tom Dieck et al., 2015) or single-molecule imaging of nascent peptides combined with single-molecule FISH, as performed in hippocampal dendrites (Wu et al., 2016).

We have shown that miR-329 and miR-495 activity and subsequent targeting of the Prr7 3’ UTR are required for Prr7 reduction in dendrites during HSD. Our findings from pLNA experiments are most consistent with an additive repressive effect of these two miRNAs on Prr7 mRNA translation. Such additive effects of multiple miRNAs binding to the same target have been demonstrated previously, for example with N-cadherin (Rago et al., 2014).

The activity-dependency of the miRNA-Prr7 interaction is evidenced by the induction of sensor activity for both miRNAs upon PTX treatment (Fig. 3h), as well as the pLNAs showing effects exclusively under stimulated conditions. However, the mechanisms leading to miR-329 and miR-495 induction appear to be different. In the case of miR-495, mature levels increase, pointing to a PTX-dependent regulation of miR-495 expression (Fig. 3i; Supplemental Fig. 10). Since this increase is preferentially observed in dendrites, it might involve increased local miRNA processing, miR-495 transport into dendrites and/or the local inhibition of miRNA degradation.

With respect to miR-329, the lack of a clear induction in mature miRNA levels suggests mechanisms at the level of the miR-329 RISC, e.g. interference of miR-329 RISC binding to the Prr7 3’UTR by an RNA-binding protein which is removed upon PTX treatment. Examples for activity-dependent miRNA-RBP interplay have been previously reported (Rajman et al., 2017; Edbauer et al., 2010; Tominaga et al., 2011; Kedde et al., 2007).

Additionally, the understanding that miR-134, miR-329 and miR-495 activity all lead to SPAR downregulation in HSD is intriguing, given that these three miRNAs are derived from the same genomic region, termed the miR-379-410 cluster located within the imprinted DLK1-DIO3 region on chromosome 14q32 in humans (da Rocha et al., 2008). Another cluster member, miR-485, also plays a role in homeostatic plasticity through expression regulation of presynaptic synaptic vesicle protein (SV2A) (Cohen et al., 2011). This shared origin of cluster miRNAs not only further support the functional significance of miR-379-410 members in activity-dependent synaptic plasticity mechanisms as previously described (Fiore et al., 2009), but also would point toward an interesting idea that individual cluster members act in distinct yet converging pathways in HSD.

### (Patho)physiological impact of the miR329/495/Prr7 pathway

A previous study (Lackinger et al., 2019) revealed that mice with a constitutive functional deletion of miR-379-410 exhibited heightened sociability and anxiety, along with increased excitatory transmission in hippocampal excitatory neurons, as well as upregulation of Prr7. These findings not only are consistent with the proposed role of Prr7 in excitatory synaptogenesis, but also point toward the connection of miRNA/Prr7 interactions to social or anxiety behavior. In other words, our current results would prompt behavioral studies examining miR-329/495/Prr7 excitatory synapse regulation in vivo. Prr7 knockout mice have been generated in previous studies with no lethal effects in the context of immune regulation (Hrdinka et al., 2016), making the study of hippocampal excitatory transmission and behavior in these mice in the context of miRNA manipulation possible.

Together with the reported involvement of Prr7 in apoptosis (Kravchick et al., 2016), our results may also suggest the importance of miRNA-dependent Prr7 downregulation in synaptic homeostasis and neuronal survival in the face of excitotoxic insult. Namely, Prr7 knockdown was shown to attenuate the excitotoxic response in hippocampal neurons following NMDAR stimulation by glutamate in a c-Jun dependent manner (Kravchick et al., 2016). Consistent with this idea, we have shown that Prr7 reduction is necessary for excitatory synapse depression upon chronic stimulation. Given our findings of miR-329 and −495-mediated Prr7 inhibition by PTX, it would therefore be reasonable to ask if these same miRNAs are activated upon glutamate stimulation, and if such activation may have neuroprotective effects against excitotoxicity. Taken together, a possible model emerges in which excessive NMDAR stimulation activates miRNAs that target Prr7, thereby reducing synapse-localized Prr7, as well as preventing Prr7 translocation to the nucleus. Consequently, the absence of dendritic Prr7 leads to spine elimination via SPAR degradation for the purpose of homeostasis, while the inhibition of nuclear Prr7 accumulation leads to c-Jun degradation for the purpose of neuronal survival.

More broadly, this idea may be tested in an in vivo context with possible future applications toward neuroprotection following status epilepticus or ischemic stroke, as NMDAR overstimulation is implicated in these conditions (McDonough and Shih, 1997; Marshall et al., 2003). Additionally, dendritic spine loss has been observed in epilepsy (Swann et al., 2000) and after stroke (Brown et al., 2007), which could indeed suggest the initiation of HSD (in addition to other neuroprotective mechanisms) to counter excitotoxicity. The therapeutic effect of miR-329/-495 administration, or Prr7 inhibition in the context of these conditions is yet to be uncovered.

## Materials and methods

### DNA constructs

All primer sequences used for cloning are indicated in Supplemental Methods.

miR-30a-chimeric hairpins for miR-329 and miR-495 stable overexpression were generated via polynucleotide cloning into the 3’ UTR of eGFP in pAAV-hSyn-EGFP vector (Addgene Plasmid #114213) using BsrGI and HindIII sites, as described previously (Christensen et al., 2010).

Control and Prr7 shRNA vectors were constructed using the pSUPER RNAi System (OligoEngine). Custom primers were designed for polynucleotide cloning into the pSUPER basic vector (VEC-PBS-0001/0002) using BglII/HindIII sites, to generate an shRNA targeting a 19-nucleotide sequence unique to rat Prr7 coding region (cggaatcggacatgtctaa).

For the Prr7 expression construct, full-length rat Prr7 coding sequence (NM_001109116.1) was amplified from hippocampal rat cDNA, then subcloned into the CMV-pcDNA3 vector using BamHI/XbaI sites. Subsequently, a start codon (atg) with HA-tag (tacccatacgacgtcccagactacgct) was inserted at the HindIII site upstream of Prr7 by polynucleotide cloning. To generate the shRNA-resistant construct, 6 point mutations in the Prr7 coding region were introduced, such that Prr7 shRNA could no longer recognize the mRNA product, yet the amino acid sequence of the resultant exogenous Prr7 protein (AESDMSK) would remain unchanged. Mutagenesis was performed using Phusion-site directed mutagenesis kit (Thermo Fisher).

Bi-cistronic reporter constructs for miRNA activity were described previously (Fiore et al., 2009). Two perfectly complementary binding sites for either miR-329 and miR-495, separated by a 2-nucleotide linker, were inserted into the dsRED 3’ UTR of the pTracer-CMV-dsRED vector, using XbaI/NotI sites by polynucleotide cloning.

Wild type and mutant Prr7 3’ UTR luciferase constructs were described and generated previously (Lackinger et al., 2018), wherein the 3’ UTR of Prr7 (NM_0010302964) was amplified from mouse DNA and cloned into the pmiRGLO dual-luciferase expression vector (Promega). Mutations in miRNA binding sites conserved across mammals were introduced by site-directed mutagenesis using Pfu Plus! DNA Polymerase (Roboklon).

### Cell culture

Primary cortical and hippocampal neuronal cultures were prepared from embryonic day 18 (E18) male and female Sprague-Dawley rats (Janvier Laboratories) as previously described (Schratt et al., 2006). Dissociated cortical neurons were directly seeded on 6-well plates coated with poly-L-ornithine (used for nucleofections), whereas hippocampal neurons were seeded on poly-L-lysine/laminin-coated coverslips in 24-well plates.

For compartmentalized cell cultures, dissociated hippocampal cells were plated onto 1-μm pore and 30-mm diameter polyethylene tetra-phthalate (PET) membrane filter inserts (Millipore) that were matrix-coated with poly-L-lysine (Sigma-Aldrich) and Laminin (BD Biosciences) on the top and bottom, also as described previously (Bicker et al., 2013; Poon et al., 2006). All neuron cultures were maintained in Neurobasal media supplemented with 2% B27, 2mM GlutaMax, 100 lg/ml streptomycin and 100U/ml penicillin (Gibco, Invitrogen) in an incubator with 5% CO_2_ at 37 °C.

HEK293T cells (Sigma) were maintained in 6 cm dishes in DMEM media containing 10% fetal bovine serum, 1mM glutamine, 100U/ml penicillin, and 100 lg/ml streptomycin (“HEK media”) in an incubator with 5% CO_2_ at 37 °C.

### Transfections and Nucleofections

All transfections of hippocampal cells were performed using Lipofectamine 2000 (Invitrogen), in triplicate wells on DIV13. 1 μg of total DNA was transfected per well in a 24-well plate, where an empty pcDNA3 vector was used to make up the total amount of DNA. Neurons were transfected in culture media in the absence of streptomycin and penicillin for 2h, replaced with neuron culture media containing ApV (1:1000) for 45min, which was washed out and replaced with conditioning media.

Nucleofections were done on cortical neurons using the P3 Primary Cell 4D-Nucleofector X Kit (Lonza, LZ-V4XP-3024), on the day of preparation and dissociation (DIV 0). 4 million dissociated cortical cells were electroporated with 3 μg total DNA per condition using the program DC-104, seeded in 6-well plates in DMEM/Glutamax supplemented with 5% FBS and incubated for 4h, then replaced with neuron culture media and incubated at 37°C until harvesting. The following amounts of DNA were used for the relevant nucleofections: 2μg chimeric miR30a-miR329 and 495 hairpins for miRNA overexpression validation; 2μg pSUPER and 1μg GFP for protein quantifications upon Prr7 knockdown; 2μg HA-Prr7 and 1μg GFP for validation of Prr7 overexpression.

HEK293T cells were transfected in HEK media supplemented with HEPES (25mM) at 1.9 million cells seeded in 6cm dishes per condition, by combining DNA with polyethylenimine (PEI) and Optimem for 15min, then adding the mixture dropwise onto cells. Cells were incubated at 37°C for 2 days until harvesting. For HA-Prr7 validation, cells were transfected with 2μg pcDNA, HA-Cav1.2, or HA-Prr7, and 1μg GFP. For shRNA-resistant mutant validation, cells were transfected with 100ng HA-Prr7 constructs, 1μg pSUPER, and 1μg GFP.

### Stimulation

To examine downscaling processes, DIV18 or DIV19 hippocampal cells were stimulated with either picrotoxin (100μM, Sigma) or equivalent volume (1:500) of ethanol absolute for 48h. For the PTX time course experiment, the picrotoxin or ethanol treatment was added in triplicate wells at DIV17 for “48h” time point, on DIV18 for “24h”, on DIV19 for “8h”, and all cells lysed together for RNA extraction 8h following the final treatment.

For investigating CDK5 inhibition, DIV19 hippocampal cells were stimulated with either Roscovitine (10μM, Sigma R7772) or equivalent volume (1:1000) of DMSO for 18h.

### Luciferase reporter assay

DIV13 primary rat hippocampal neurons were transfected in triplicate with 20pmol pLNAs (10pmol for 329/495 mix) and 50ng Prr7 3’ UTR pmiRGLO constructs per well. Cells were treated with PTX or ethanol on DIV18 or DIV19 for 48h, then lysed in Passive Lysis Buffer (diluted to 1x, Promega) for 15min, and dual-luciferase assay performed using homemade reagents (as described in Baker et al., 2014) on the GloMax Discover GM3000 (Promega).

pLNAs used were: Control pLNA (miRCURY LNA miRNA Power Inhibitor Negative control A, Qiagen #339135 YI00199006-DCA), miR-329 pLNA (miRCURY LNA miRNA Power Inhibitor RNO-MIR-329-3P, Qiagen # 339130 YI04101481-DCA), miR-495 pLNA (miRCURY LNA miRNA Power Inhibitor HSA-MIR-495-3P, Qiagen #339130 YI04101229-DCA).

### Bicistronic reporter (dual color) assay/single cell fluorescent sensor assay

For single cell fluorescent assay, DIV13 or DIV14 hippocampal cells were transfected in triplicate wells with 125ng of control, miR-329, or miR495 bicistronic sensor, and 5pmol control, miR-329, or miR-495 pLNA (where applicable). On DIV19, cells were treated with PTX or ethanol for 48h, then fixed for 15min in 4%paraformaldehyde/4% sucrose/PBS, washed in PBS and directly mounted onto slides for imaging. To determine miRNA activity, dsRED-positive (red or yellow) vs. GFP-only (green) cells were manually counted at 20x objective with both 488 and 561 channels open for all coverslips, and the proportion of GFP-only cells over the total count was taken. Approximately 100-200 cells were counted per coverslip (300-600 total per experimental condition).

### Immunocytochemistry, spine density and image analysis

For all imaging experiments, hippocampal cells were transfected on DIV13 in either duplicate or triplicate wells. The following amounts of DNA/RNA were used for the relevant experiments: 150ng GFP-amp, 20pmol pLNAs (10pmol for 329/495 mix), 500ng miR30a-hairpins, 7.5ng pSUPER constructs (with the exception of spine rescue experiment with pLNAs, for which 2.5ng was used), 400ng HA-Prr7 or HA-Prr7R. Where applicable, the transfected cells were further treated on DIV19 with picrotoxin/ethanol or roscovitine/DMSO.

For all experiments, cells were fixed for 15min with 4% paraformaldehyde/4% sucrose/PBS and washed with PBS. In cases where cells were only analyzed for spine morphology, coverslips were directly mounted onto microscope slides.

For immunostaining, following fixation coverslips were transferred to a humidified chamber protected from light. For Prr7 immunostaining only, cells were permeabilized with 0.1% Triton/PBS for 5min. Blocking for 30min in 1xGDB buffer (0.02% gelatin/0.5% Triton X-100/PBS) was followed by overnight incubation with primary antibody in GDB at 4°C. Secondary antibodies in GDB were applied for 45min. Coverslips were washed with PBS before and after fixation, and application of each antibody, and briefly in MilliQ before slide mounting. The following primary antibodies were used: Rabbit polyclonal anti-Prr7 (200ng/ml, PA5-61266, Invitrogen), Chicken monoclonal anti-Map2 (1:5000 dilution, PA1-16751, Thermo Fisher), rabbit polyclonal anti-SPAR (1:1500 dilution, kind gift of D.T. Pak). Alexa-546 and - 647-conjugated secondary antibodies (1:20,000 dilution) were used for detection.

All images were acquired with confocal laser-scanning microscope (Zeiss LSM) using a 40x/1.3 oil DIC UV-IR M27 objective. Z-stack images were obtained for 7-11 GFP-positive neurons with pyramidal morphology for each condition. Settings were: Opt sampling (1.0x Nyquist), Zoom factor 1.0, Pixel Dwell 0.90μs, Speed fps 0.23, Scan time 4.32s, Speed 6, Digital Gain 1.0, Pinhole 384μm. Z stacks were kept at 0.45μm and 9-11 slices obtained. Laser settings were kept constant between conditions. Images were processed by Airyscan processing at 6.0 strength 3D, and maximum intensity projections of the Z-stacks were used for signal quantification.

Prr7 and SPAR puncta intensities were analyzed with a custom-made Python-script developed by D. Colameo and can be added as a Plugin on Fiji (https://github.com/dcolam/Cluster-Analysis-Plugin). Whole cell, cell body, and dendrites (whole cell selection with cell body subtracted) were defined using GFP as a mask.

Spine density was measured manually using Fiji. For each cell analyzed, first a primary dendrite and one secondary dendrite branching off it was selected, and the total length of the selected segment obtained. Using the “cell counter” tool, the total spine number along the selected segment was counted (without discrimination of spine shape), and the count was divided by the total length to obtain #spines/μm. A second dendritic segment was selected, and the process repeated. The average of the two spine density readings was calculated per cell.

### Preparation of protein extracts and western blotting

Protein extracts were prepared by first scraping and lysing cells in RIPA buffer (150mM NaCl, 1% Triton X-100, 0.5% Sodium Deoxycholate, 1mM EDTA, 1mM EGTA, 0.05% SDS, 50mM Tris pH 8.0, 1x complete protease inhibitor cocktail (Roche)), spinning down the lysate at maximum speed at 4°C for 15min, and collecting the supernatant. Protein concentration was measured using the Pierce BCA Protein Assay Kit (Thermo Fisher). Equal amounts of protein were diluted in Laemmli sample buffer supplemented with BME, boiled at 95°C for 5min and loaded onto SDS-PAGE gels (10% polyacrylamide for Prr7 probe, 8% for SPAR). For Prr7 western, proteins were transferred onto Trans-Blot Turbo 0.2μm nitrocellulose membranes (Biorad) using the Trans-Blot Turbo semi-dry transfer system (Biorad). For SPAR western, proteins were blotted onto 0.45μm PVDF membranes (Immobilon) soaked in transfer buffer (25mM Tris-HCl pH 8.3, 192mM Glycine, 20% MeOH) via wet transfer for 15-16h at 25V. For all experiments, blocking was done in 5% milk in 1xTBS-0.1% Tween20 (TBST) for 1.5-2h at room temperature, followed by overnight primary antibody incubation at 4°C. Following washes in milk, horse radish peroxidase (HRP)-conjugated secondary antibodies were applied onto the membranes for 1h at room temperature. Membranes were washed in TBST and visualized with the Clarity Western ECL Substrate (Biorad) on the ChemiDoc Imaging System (Biorad). The following primary antibodies were used: Rabbit anti-GluA1 (1:1000 dilution, PA1 37776, Thermo Fisher), Mouse monoclonal anti-Prr7 (1:250 dilution, MA1-10448, Thermo Fisher), Rabbit monoclonal anti-alpha Tubulin (1:2000 dilution, 11H10 lot 112125S, Cell signaling). HRP-conjugated secondary antibodies Rabbit anti-Ms IgG H&L (402335 lot D00160409, Calbiochem), Goat anti-Rb IgG H&L (lot 2625715, Calbiochem) were used at 1:20,000 dilution.

### RNA extraction and Quantitative real-time PCR

RNA was isolated using TriFast RNA extraction kit (30-2030, VWR) or RNA-Solv reagent (Omega Bio-tek). Genomic DNA was removed with TURBO DNAse enzyme (Thermo Fisher). Reverse transcription was performed using either the Taqman MicroRNA Reverse Transcription Kit (Thermo Fisher) for miRNA detection, or iScript cDNA synthesis kit (Biorad) for mRNA detection. qPCR was performed using either Taqman Universal PCR Master Mix (Thermo Fisher) for microRNA detection, or the iTaq SYBR Green Supermix with ROX (Biorad), and plates were read on the CFX384 Real-Time System (Biorad). Data was analyzed via ΔΔCt method, and normalized to either U6 (for miRNAs) or GAPDH (for mRNAs). mRNA primer information is indicated in supplemental methods.

Taqman primers used were (all from Thermo Fisher): U6 snRNA (Assay ID: 001973), mmu-miR-495 (4427975, Assay ID: 001663), mmu-miR-329 (4427975, Assay ID: 00192), mmu-miR-134 (4427975, Assay ID: 001186), hsa-miR-132 (4427975, Assay ID: 000457), hsa-miR-99b-5p (4427975, Assay ID: 000436).

### Statistics

Statistical tests were performed using GraphPad Prism version 9.2.0. For all datasets, three to four independent experiments were performed. Given the small sample size, normality was assumed for all datasets, and therefore one or two sample Student’s t-test (two-sided), or oneway or two-way ANOVA followed by post-hoc Tukey test were performed. * P < 0.05; ** P < 0.01; *** P < 0.001.

## Acknowledgements

We would like to thank D. T. Pak and M. Sheng for generously providing the SPAR antibody. We thank M. Soutschek, R. Fiore, S. Bicker, C. Gilardi for cloning the pmiRGLO, miR-329/control sensors, Control pSUPER, and miR30a-Ctr hairpin constructs respectively. We thank the excellent technical assistance of T. Wüst and Cristina Furler. This work was supported by a grant from the Swiss National Foundation (SNF 310030_205064) to G.S.

## Author Contributions

MOI performed all experiments and analyses except for compartmentalized cell culture (plasmid cloning, transfections, luciferase assays, immunocytochemistry, western blotting, imaging, RT-qPCR), and wrote the manuscript. DC performed compartmentalized RNA/protein extractions and wrote the Python script for puncta analysis. IA contributed qPCR data. GS conceptualized and supervised the project, and wrote the manuscript.

