## Supplemental Material for "miR-329 and miR-495-mediated Prr7 downregulation is required for homeostatic synaptic depression in rat hippocampal neurons"

**Supplemental Methods:**

**Primer sequences (all 5’ to 3’)**

miR-329 hairpin:

Oligo A-329-BsrGI for: GTACAGCTGTTGACAGTGAGCGACAACACACCC

Oligo A-329-BsrGI rev: AGGTTAGCTGGGTGTGTTGTCGCTCACTGTCAACAGCT

Oligo B-329 for: AGCTAACCTTTTTTGTGAAGCCACAGATGGAAAAAGGT

Oligo B-329 rev: CCCAGAACCTTTTTCCATCTGTGGCTTCACAAAAA

Oligo C-329-HindIII for: TCTGGGTGTGTTGCTGCCTACTGCCTCGGAA

Oligo C-329-HindIII rev: AGCTTTCCGAGGCAGTAGGCAGCAACACA

miR-495 hairpin:

Oligo A-495-BsrGI for: GTACAGCTGTTGACAGTGAGCGACAAACAAACA

Oligo A-495-BsrGI rev: AGTGCACCATGTTTGTTTGTCGCTCACTGTCAACAGCT

Oligo B-495 for: TGGTGCACTTCTTTGTGAAGCCACAGATGGAAGAAGTG

Oligo B-495 rev: ACATGGCACTTCTTCCATCTGTGGCTTCACAAAGA

Oligo C-495-HindIII for: CCATGTTTGTTTGCTGCCTACTGCCTCGGAA

Oligo C-495-HindIII rev: AGCTTTCCGAGGCAGTAGGCAGCAAACAA

Control hairpin:

Oligo A-Ctr-BsrGI for: TGTACAGCTGTTGACAGTGAGCGACAACCTTGTG

Oligo A-Ctr-BsrGI rev: AAGGACCACAAGGTTGTCGCTCACTGTCAACAGC

Oligo B-Ctr for: GTCCTTAGGTGCGTGTGAAGCCACAGATGGCGC

Oligo B-Ctr rev: GGTTTAGGTGCGCCATCTGTGGCTTCACACGCACCT

Oligo C-Ctr-HindIII for: ACCTAAACCACAAGGTTGCTGCCTACTGCCTCGGA

Oligo C-Ctr-HindIII rev: AAGCTTTCCGAGGCAGTAGGCAGCAACCTTG

pSUPER:

Prr7 shRNA targeting rat (and mouse) Prr7 coding region cggaatcggacatgtctaa:

siPrr7_for 1: GATCCCC CGG AAU CGG ACA UGU CUA A TTCAAGAGA

siPrr7_rev 1: AGCTTAAAAA CGG AAU CGG ACA UGU CUA A TCTCTTGAA

siPrr7_for 2: UUA GAC AUG UCC GAU UCC G TTTTTA

siPrr7_rev 2: UUA GAC AUG UCC GAU UCC G GGG

Control shRNA:

siCtr_for1: GATCCCCAAACCTTGTGGTCCTTAGGTTCAAGAGA

siCtr_rev1: AGCTTAAAAAAACCTTGTGGTCCTTAGGTCTCTTGAA

siCtr_for2: CCTAAGGACCACAAGGTTTTTTTA

siCtr_rev2: CCTAAGGACCACAAGGTTTGGG

HA-Prr7:

Amplification of rat Prr7 coding sequence with BamHI and XbaI sites:

For:  TATAGGATCCGTGATGTCCCAGGGCA

Rev: TCACTCTAGACTATACGGCTGTAGTCCTCCC

Start codon and HA-tag insertion using HindIII site:

For: AGCTT ATG TACCCATACGACGTCCCAGACTACGCT A

Rev: AGCTT AGCGTAGTCTGGGACGTCGTATGGGTA CAT A

shRNA-resistant HA-Prr7:

For: CGG CCC TGG AGC TAT CCG CGC CAA gCC GAG TCA GAT ATG AGT AAg CCG CCG TGC TAC GAG GAG GCG GTG

Rev: CAC CGC CTC CTC GTA GCA CGG CGG cTT ACT CAT ATC TGA CTC GGc TTG GCG CGG ATA GCT CCA GGG CCG

miRNA sensor:

miR-495 mature sequence: AAACAAACAUGGUGCACUUCUU

miR-495 for: GGCCGC aagaagtgcaccatgtttgttt ca aagaagtgcaccatgtttgttt T

miR-495 rev: CTAGA aaacaaacatggtgcacttctt tg aaacaaacatggtgcacttctt GC

(used in Fiore et al., 2009)

miR-329 mature sequence: aacacacccagcuaaccuuuuu

miR-329 for : ggccgc-aaaaaggttagctgggtgtgtt-AC-aaaaaggttagctgggtgtgtt-T

miR-329 rev : ctaga-aacacacccagctaacctttt-GT-aacacacccagctaacctttt-gc

miR-Ctr for :
GGCCGCAAGGGATTCTGATGTTGGTCACACTACAAGGGATTCTGATGTTGGTCACACTT

miR-Ctr rev :
CTAGAAGTGTGACCAACATCAGAATCCCTTGTAGTGTGACCAACATCAGAATCCCTTGC

(used in Lackinger et al., 2018)

Prr7 3’ UTR luciferase reporter

Prr7 3’ UTR for: AAACTCGAGAGGACTACAGCCGTATAGAGG

Prr7 3’ UTR rev: TTTGTCGACGTACCAAAGCAGATCACACACC

Prr7 3’ UTR mutagenesis was performed for each miRNA binding site sequentially.

Prr7 3’ UTR mut1 for: TACCCTGTTGAATTCATTTTGAGGATAATAAAGG

Prr7 3’ UTR mut1 rev: TCCTCAAAATGAATTCAACAGGGTAAGAAATCC

Prr7 3’ UTR mut2 for: ATAATAAAGGTCTAGAATCTGCTTTGGTACGtCG

Prr7 3’ UTR mut2 rev: ACCAAAGCAGATTCTAGACCTTTATTATCCTCAAAATG

**qPCR primers**

GAPDH for: GCCTTCTCTTGTGACAAAGTGGA

GAPDH rev: CCGTGGGTAGAGTCATACTGGAA

Prr7 for: GTCACGCCCTTTCTGAGC

Prr7 rev: ATGCAGCGCCGAGGTATA

GluA1 for: CGAGTTCTGCTACAAATCCCG

GluA1 rev: TGTCCGTATGGCTTCATTGATG

cFos for: CATCATCTAGGCCCAGTGGC

cFos rev: AGGAACCAGACAGGTCCACATCT

### Supplemental Figure Legends

**Figure 1 Prr7 polyclonal antibody validation for immunostaining. (a)** Titration series testing concentrations 10-800ng/ml of antibody by quantifying cell body Prr7 signal intensity in 7 neurons transfected with Ctr vs. Prr7 shRNA. DIV12 rat hippocampal cells were transfected with 150ng GFP-Amp plasmid, 7.5ng pSUPER control or Prr7 shRNA vector with pcDNA added up to 1µg total per well. On DIV18 cells were fixed 15min with 4% PFA/4% sucrose/PBS, blocked in 1xGDB for 15min, and immunostained for Prr7 (PA5-61266 Thermo at given concentrations) in GDB overnight at 4C. The following day coverslips were washed, secondary (546 anti-Rb for Prr7, 1:2000) added and 7-9 neurons imaged. ** $p < 0.01$, *** $p < 0.001$, **** $p < 0.0001$, 2-way ANOVA with Tukey's post-hoc HSD test. **(b)** Representative images of Ctr and Prr7 shRNA-transfected neurons stained for Prr7 using 200ng/ml antibody, with close-ups of dendrites and cell bodies. Arrows indicate that some, but not all, of dendritic spines colocalize with Prr7 puncta. **(c)** Results of average Prr7 punctum intensity in processes (cell body puncta subtracted) of imaged cells (*** $p=0.0007$, unpaired Student's t-test).

**Figure 2 Segmentation analysis of cells treated with EtOH or PTX for 48h and immunostained for Prr7. (a)** The same EtOH and PTX-treated cells as those analyzed for whole cell, cell body, dendrite Prr7 levels in Figure 1g-h were re-visited to measure Prr7 immunofluorescence with respect to distance from the soma. Through an automated method, concentric circles increasing in 10µm steps were drawn around the soma, and dendritic segments within each increasing step were selected using GFP as a mask. Subsequently the average Prr7 puncta intensity in the detected dendritic segments (as measured by integrated density) were obtained, and a grand average across the 7-9 cells imaged per condition was calculated for one experiment. Shown is the mean Prr7 immunofluorescence data across 3 independent experiments, normalized to the Mock 10µm point, ± SEM. **(b)** A sample image showing dendritic segments selection and Prr7 puncta detected at 50µm from the soma through the automated analysis.

**Figure 3 Raw pre-normalized Prr7 mRNA and protein data from compartmentalized experiments.** The same data from Fig. 1b, e, g are shown, but before normalization to the EtOH-treated conditions. Namely, **(a)** represents Prr7 mRNA levels in compartmentalized hippocampal cell samples treated at DIV19 with 100µM PTX or EtOH (1:500 volume) for 48h. **(b)** Prr7 protein levels as measured by mean optical band density relative to Tubulin in compartmentalized hippocampal rat cultures treated at DIV19 with PTX or EtOH for 48h. Mock Cell Body vs. PTX Cell Body: ns $p=0.3612$; Mock Dendrites vs. PTX Dendrites: * $p=0.0495$; 2-way ANOVA with Tukey's post-hoc test. **(c)** Average Prr7 punctum intensity in GFP-transfected (150ng) cell body or dendrites selection of hippocampal rat cultures treated with PTX or EtOH on DIV19 for 48h, then immunstained for Prr7. For all graphs, data = mean ± S.D. , n=3-4.

**Figure 4 Prr7 protein levels in Ctr shRNA and Prr7 shRNA-nucleofected cells (validation of Prr7 knockdown with pSUPER construct).** Prr7 protein levels as measured by mean optical band density relative to Tubulin in empty pSUPER, Ctr shRNA, and Prr7 shRNA (2µg)-nucleofected cortical neuron whole cell extracts, with corresponding western blot images (membranes were cut horizontally to probe for Prr7 (29kD), GluA1 (101kD), and Tubulin (50kD)). Protein extracts were loaded in the order of pSUPER empty -> Ctr shRNA -> Prr7 shRNA. The GluA1 data is presented in Fig. 2g-h. Data = mean normalized to pSUPER empty condition ± S.D., n=3, **** $p<0.0001$, unpaired Student's t-test.

**Figure 5 Validations of Prr7 overexpression with HA-Prr7 construct. (a)** HEK293 cells were transfected with pcDNA, HA-Cav1.2, or HA-Prr7 (HA-tag fused to N-terminus of Prr7 cDNA sequence) (all 2µg) and 1µg GFP (for visualization of transfection efficiency), then protein extracts from the cell lysate were immunoblotted for HA (left) and Prr7 (right) detection. Although the HA tag of the HA-Cav1.2 was not detected, this may be due to inadequate transfer given the large size of Cav1.2 (expect a band at 250kD). However, strong bands for HA and Prr7 were detected at the expected molecular weight of 29kD for the HA-Prr7 condition only, suggesting Prr7 overexpression in HEK cells which do not express endogenous Prr7. **(b)** Prr7 protein levels as measured by mean optical band density relative to Tubulin in 2µg pcDNA or HA-Prr7 with 1µg GFP-nucleofected cortical neuron protein extracts, with corresponding western blot images (membranes were cut horizontally to probe for for Prr7 (29kD), GluA1 (101kD), and Tubulin (50kD)). Data = mean ± S.D., n=3, $p=0.0587$, unpaired Student's t-test. **(c)** Representative images of GFP (left) and Prr7 staining (right) in hippocampal rat neurons transfected with 400ng pcDNA or HA-Prr7 and 150ng GFP at DIV13, fixed at DIV18 and immunostained for Prr7. Importantly, the Prr7 overexpression appeared in not only the cell body but also in dendrites of HA-Prr7 transfected cells.

**Figure 6 Validation of shRNA-resistant Prr7 expression construct. (a)** Position of 6 point mutations in Prr7 cDNA sequence to generate shRNA-resistant construct, using generated HA-Prr7 construct as a template. Prr7 shRNA targeting region is in bold text. The mutations introduced interfere with the recognition of Prr7 mRNA by shRNA, but the corresponding amino acid sequence of the resultant exogenous Prr7 protein (AESDMSK) remains unchanged. **(b)** HEK293 cells were transfected with 100ng of either HA-Prr7 wild type (HA-Prr7 wt) or shRNA-resistant mutant (HA-Prr7$^{R}$), 1µg Ctr or Prr7 shRNA, and 1µg GFP (for assessing transfection efficiency). Protein extracts were obtained and immunoblotted for Prr7 and Tubulin expression. Prr7 knockdown due to Prr7 shRNA transfection is prevented in the presence of the mutant, but not the wild type expression construct.

**Figure 7 Sample images from miRNA sensor assay.** Representative images of miR-329, miR-495, Control sensor-transfected hippocampal neurons (DIV13 transfection, DIV19 EtOH or PTX treatment, DIV21 fixation). GFP (left), dsRed (middle), and merged (right) images are shown. Cells that appear red or yellow in the merged channel were counted as "miRNA-negative" and those that appear green were counted as "miRNA-positive".

**Figure 8 Construction of miR-329 and miR-495 hairpins for overexpression. (a)** Secondary structures and sequence of the miR30a-329-3p and miR-30a-495-3p chimeric hairpins. Shaded regions indicate the respective mature miRNA sequences expressed in the chimeric hairpins. **(b)** Expression of miR-134 (an unrelated miRNA, to assess specificity of the two hairpins), miR-329, and miR-495 assessed via qPCR and normalized to U6 levels, in primary cortical rat neurons nucleofected with Ctr, miR-495, or miR-329 hairpin (2µg AAV). All data were further normalized to the Ctr hairpin condition. As expected, miR-134 were unaffected by miR-329 and miR-495 hairpin-nucleofected cells. Specific, but moderate (~2x from control) increases in miR-329 and miR-495 levels were observed for the miR-329 and miR-495 hairpin-nucleofected cells respectively.

**Figure 9 Dendritic Prr7 protein levels show decreasing trends in miR-495 and miR-329 hairpin transfected hippocampal neurons.** Average Prr7 punctum intensity in dendrite selections of **(a)** miR-495 hairpin and **(b)** miR-329 hairpin-transfected (500ng hairpin, DIV13) hippocampal rat neurons, which were fixed on either DIV21 (miR-495) or DIV18/19 (miR-329). Puncta intensity was measured by either mean value (left bar graphs) or integrated density (right bar graphs). Data = mean ± S.D., n=3, paired Student's t-test. 495 hairpin: p= 0.0922 (left), p=0.2987 (right). 329 hairpin: p=0.2610 (left), p=0.0996 (right). Scale bars = 5µm.

**Figure 10 Whole Cell expression levels of mature miRNAs of interest in 8, 24, 48h PTX-treated hippocampal cells.** Hippocampal neurons were treated with 100µM PTX or equivalent volume of ethanol at DIV17 for 48h time point for three wells respectively, then at the same time of day on DIV18, these treatments were repeated for another three wells for the 24h time point. In the morning of DIV19, the 8h time point treatments were added, and in the evening of DIV19, cells were lysed and assessed by qPCR for mature miRNA levels. miR-99b and miR-132 were used as negative and positive controls respectively. Data = mean normalized to EtOH condition ± S.D., n = 3.

**Figure 11 Segmentation analysis of cells transfected with Control or Prr7 shRNA, treated with EtOH or PTX, and immunostained for SPAR.** The same cells as those analyzed for dendrite SPAR levels in Figure 5c were re-visited to measure SPAR immunofluorescence with respect to distance from the soma. Through an automated method, concentric circles increasing in 10μm steps were drawn around the soma, and dendritic segments within each increasing step were selected using GFP as a mask. Subsequently the average SPAR puncta intensity in the detected dendritic segments (as measured by integrated density) were obtained, and a grand average across the 7-10 cells imaged per condition was calculated for one experiment. Shown is the mean SPAR immunofluorescence data across 3 independent experiments, normalized to the Mock 10μm point, ± SEM.

### Supplemental Figures

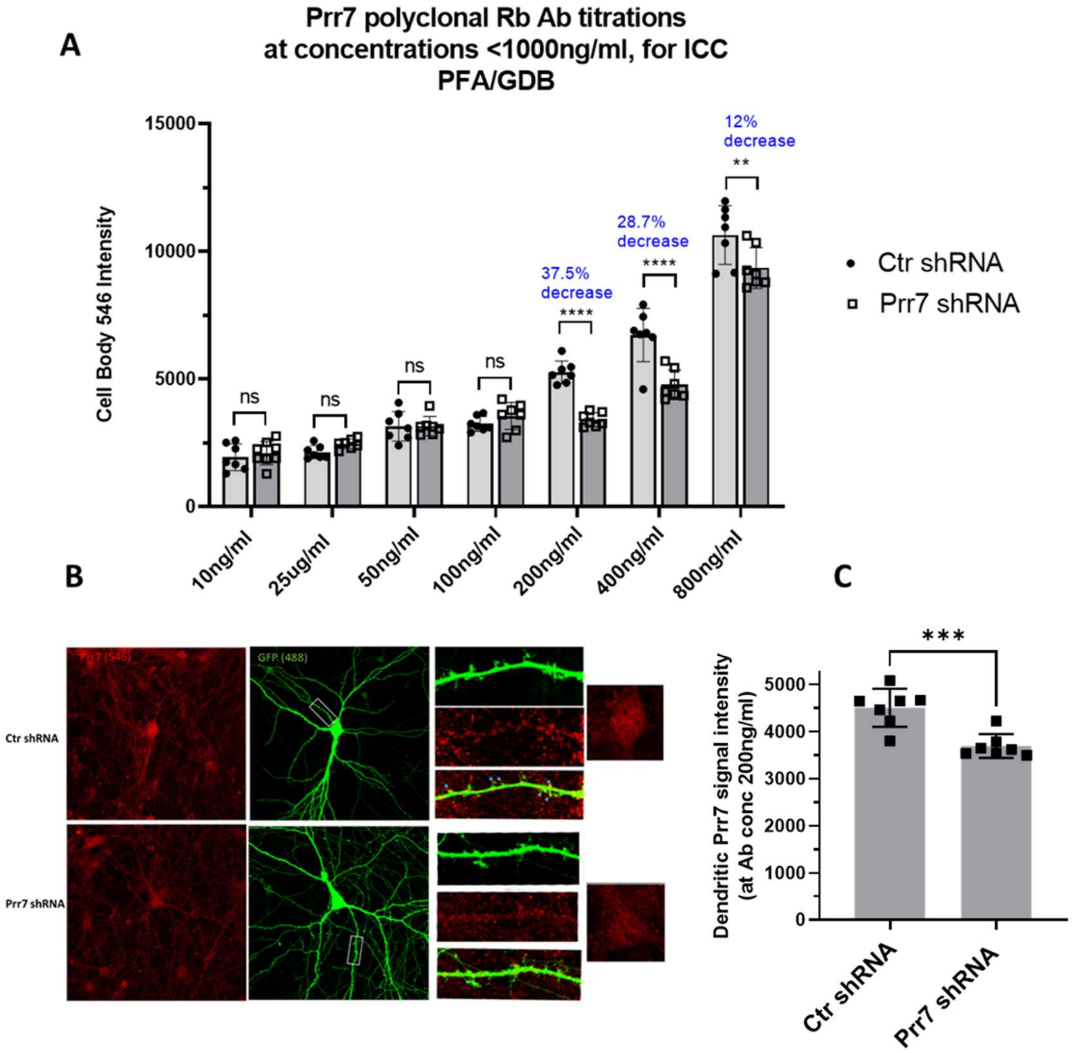


### Supplemental Figure 1.

A

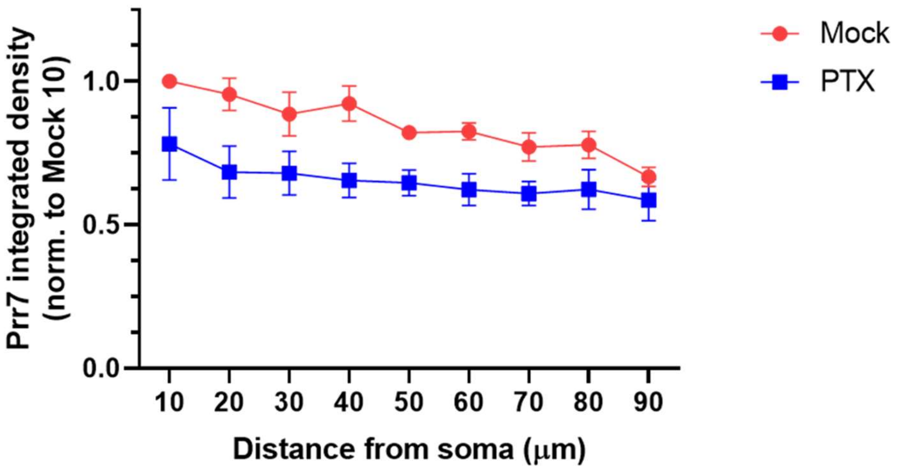


B

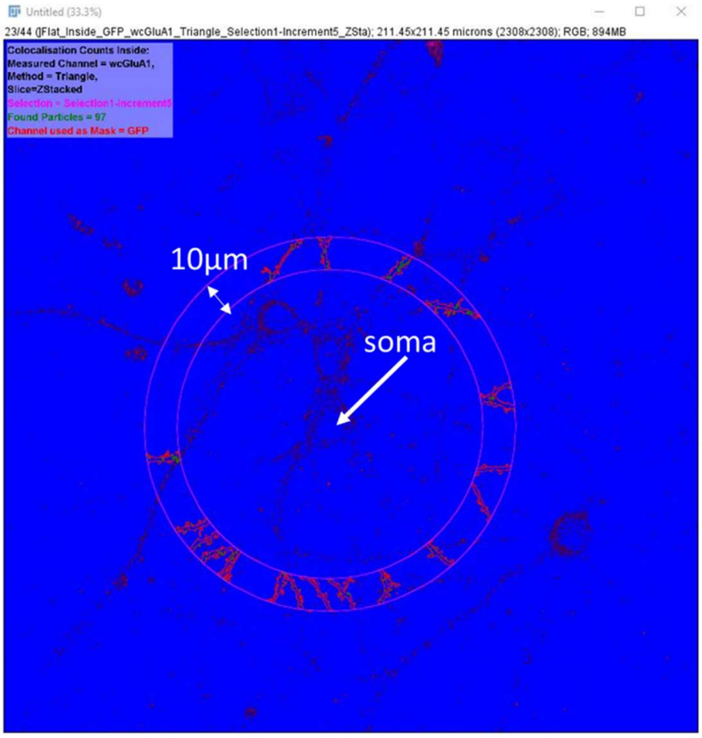


#### Supplemental Figure 2.

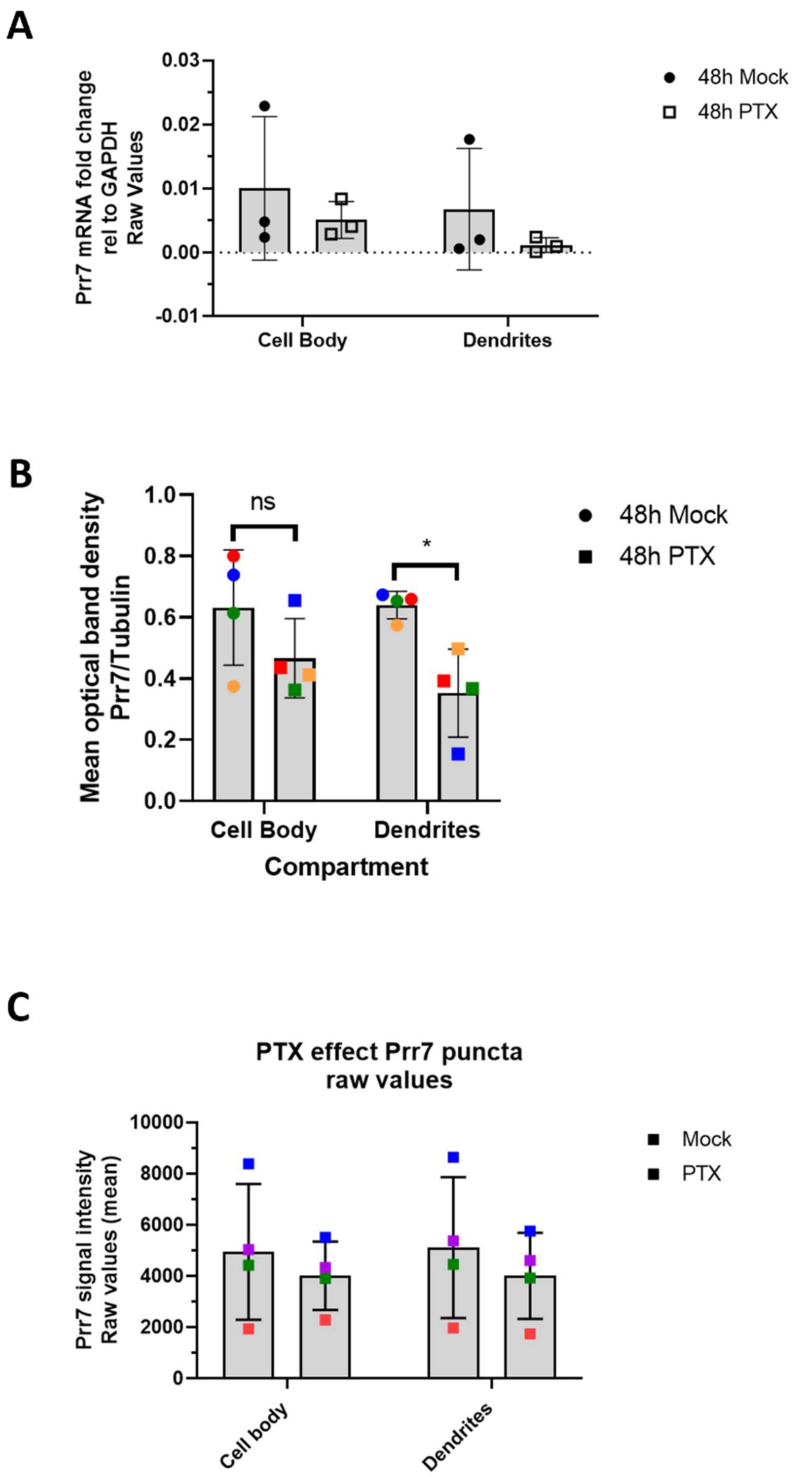


**Supplemental Figure 3.**

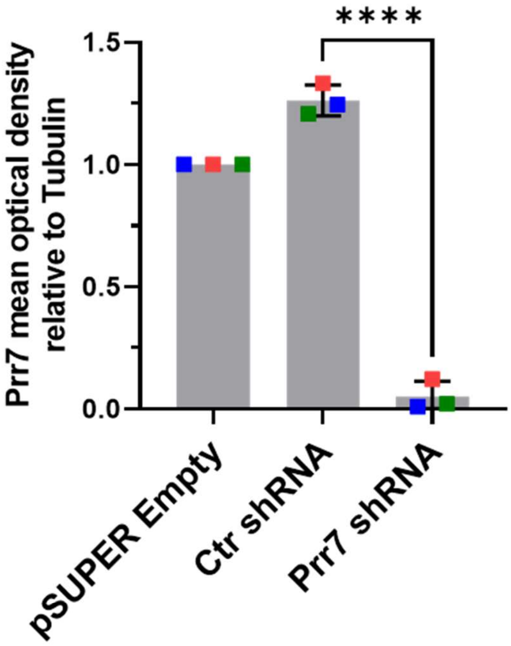


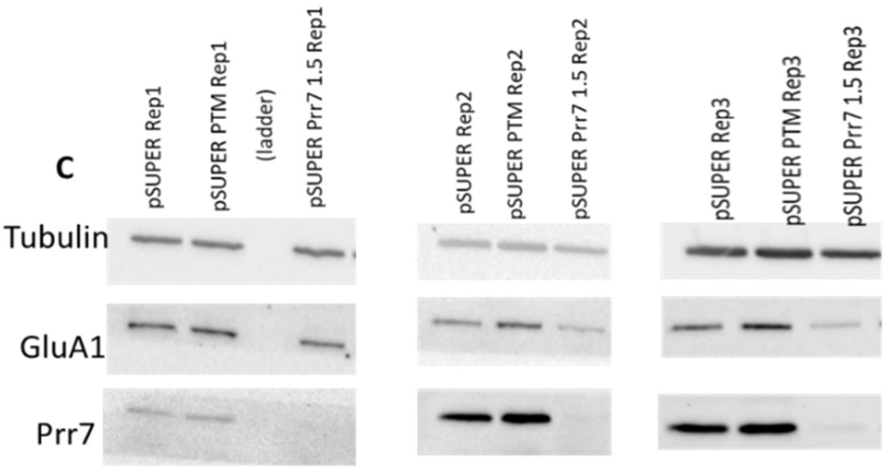


**Supplemental Figure 4.**

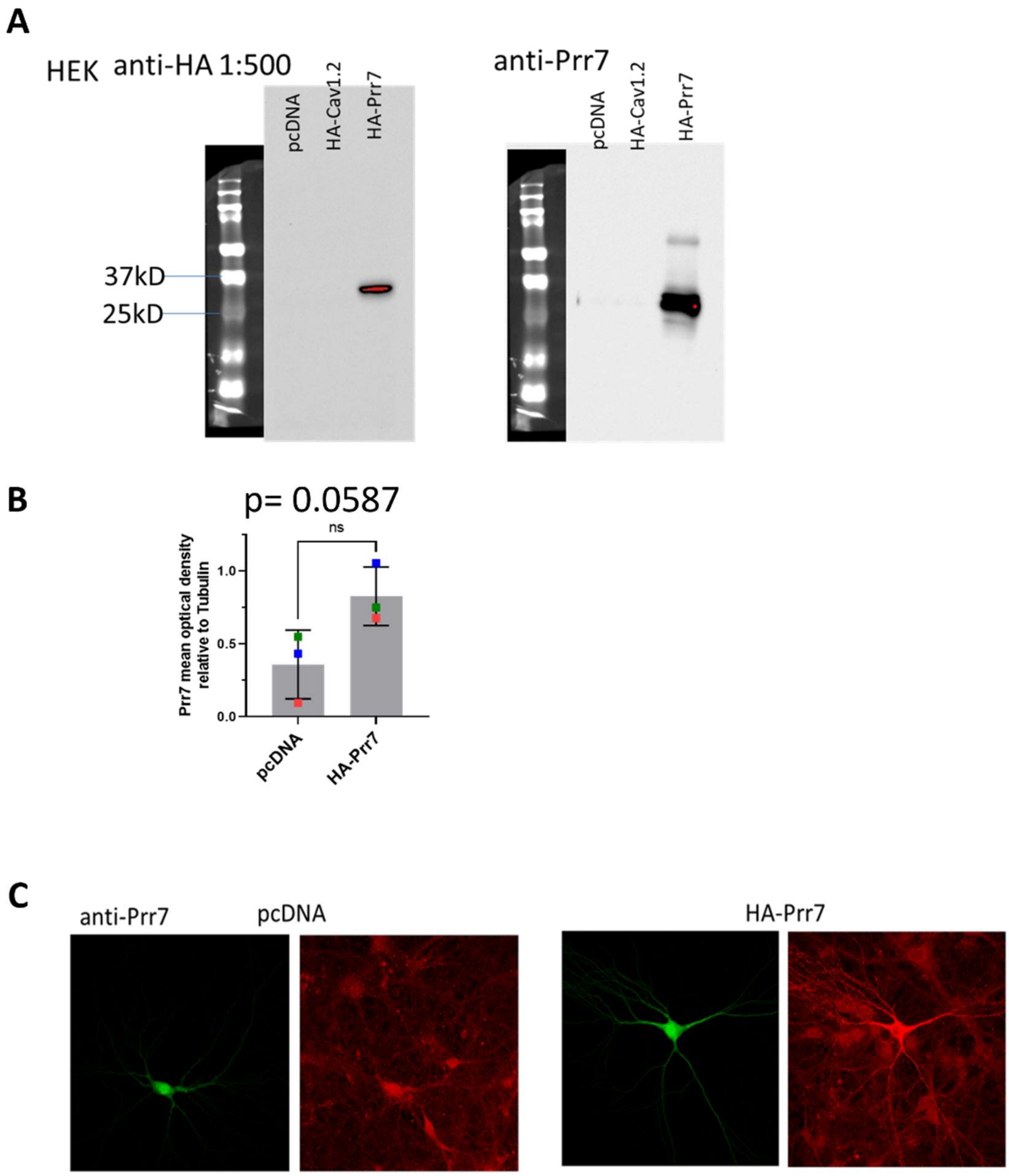


**Supplemental Figure 5.**

**A**

```
        Y   P   R   Q   A   E   S   D   M   S   K   P   P   C   Y
Fnor C TAT CCG CGC CAA gCG GAA TCG GAC ATG TCT AAg CCG CCG TGC TAC G
Rnor G ATA GGC GCG GTT cGC CTT AGC CTG TAC AGA TTc GGC GGC ACG ATG C

        Y   P   R   Q   A   E   S   D   M   S   K   P   P   C   Y
Fmut C TAT CCG CGC CAA gCC GAG TCA GAT ATG AGT AAg CCG CCG TGC TAC G
Rmut G ATA GGC GCG GTT cGG CTC AGT CTA TAC TCA TTc GGC GGC ACG ATG C
```

**B**

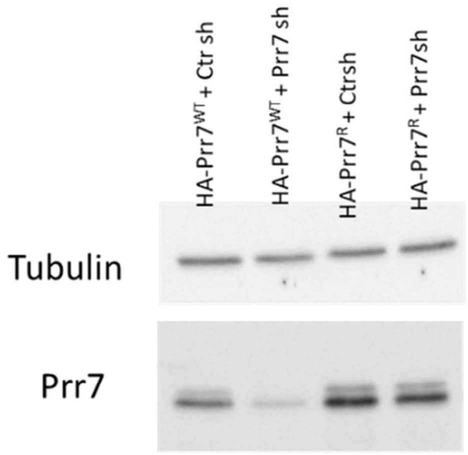


**Supplemental Figure 6.**

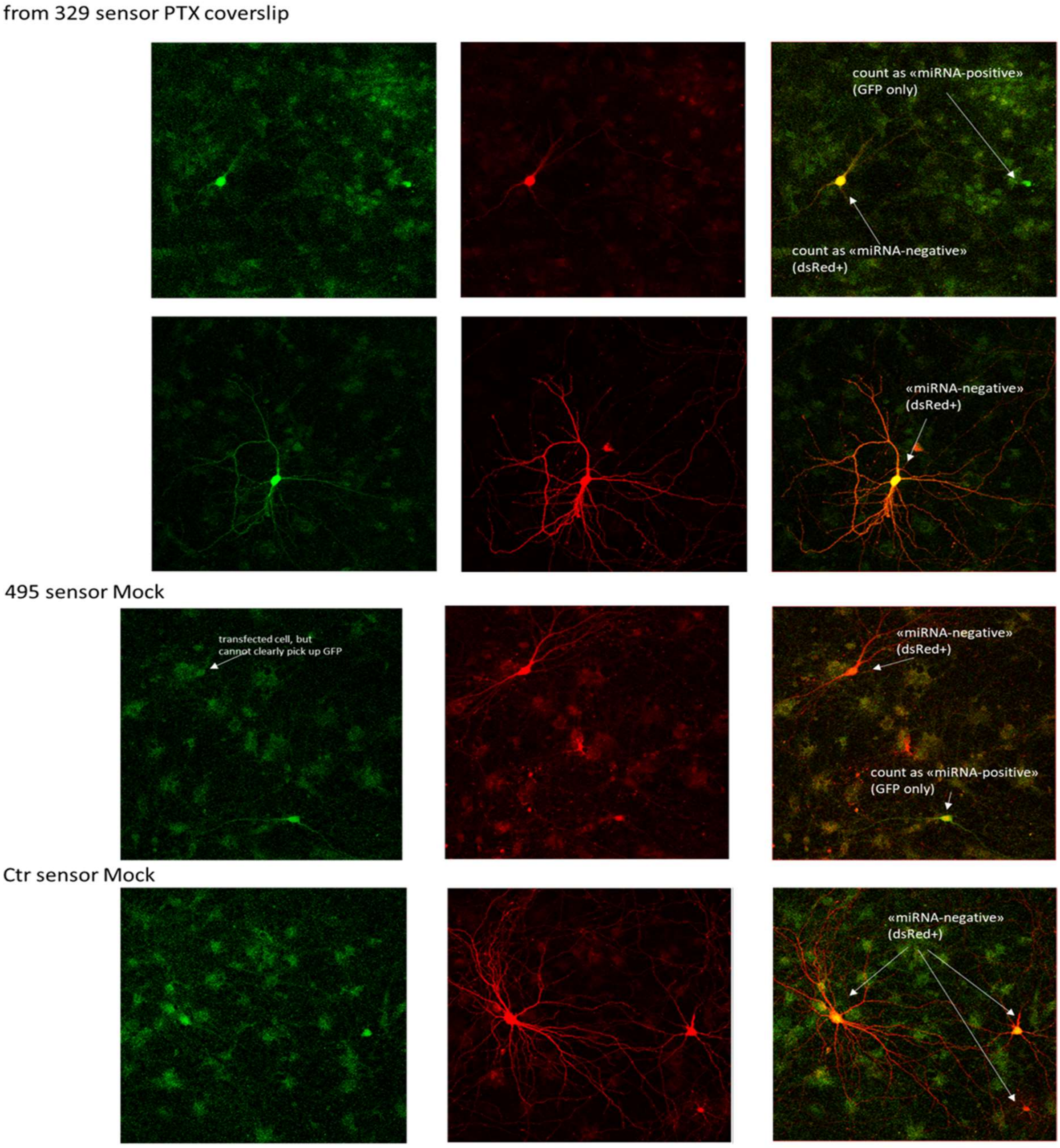


### Supplemental Figure 7.

**A**

**miR30a-329-3p oe hairpin**

>rno-miR-329-3p MIMAT0000566
AACACACCCAGCUAACCUUUUU

Note that the mmu and rno miR-329-3p mature sequences are the same.

```
-g    uuga     a   a           cu          -----   a
5' cug    cagug gcg caacacacccag  aaccuuuuuu     gug a
   |||    ||||| ||| ||||||||||||  ||||||||||     |||
3' ggc    gucau cgu guugugugggguc  uuggaaaaag     cac g
 -a   ucc-     c   c           --          guaga   c
```

**miR30a-495-3p oe hairpin**

>rno-miR-495 MIMAT0005320
AAACAAACAUGGUGCACUUCUU

miR-495-3p mature sequence is conserved for human, mouse and rat.

```
-g    uuga     a   a           gu          -----   a
5' cug    cagug gcg caaacaaacaug  gcacuucuuu     gug a
   |||    ||||| ||| ||||||||||||  ||||||||||     |||
3' ggc    gucau cgu guuuguuuguac  cgugaagaag     cac g
 -a   ucc-     c   c           --          guaga   c
```

**B**

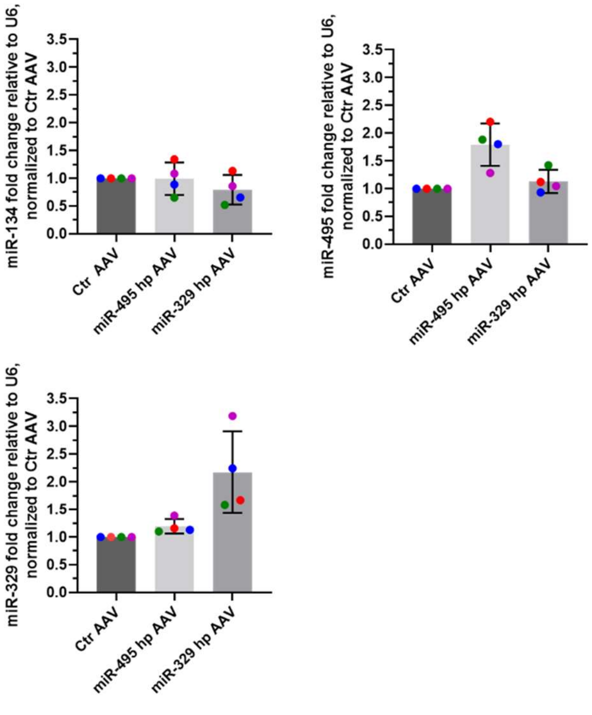


### Supplemental Figure 8.

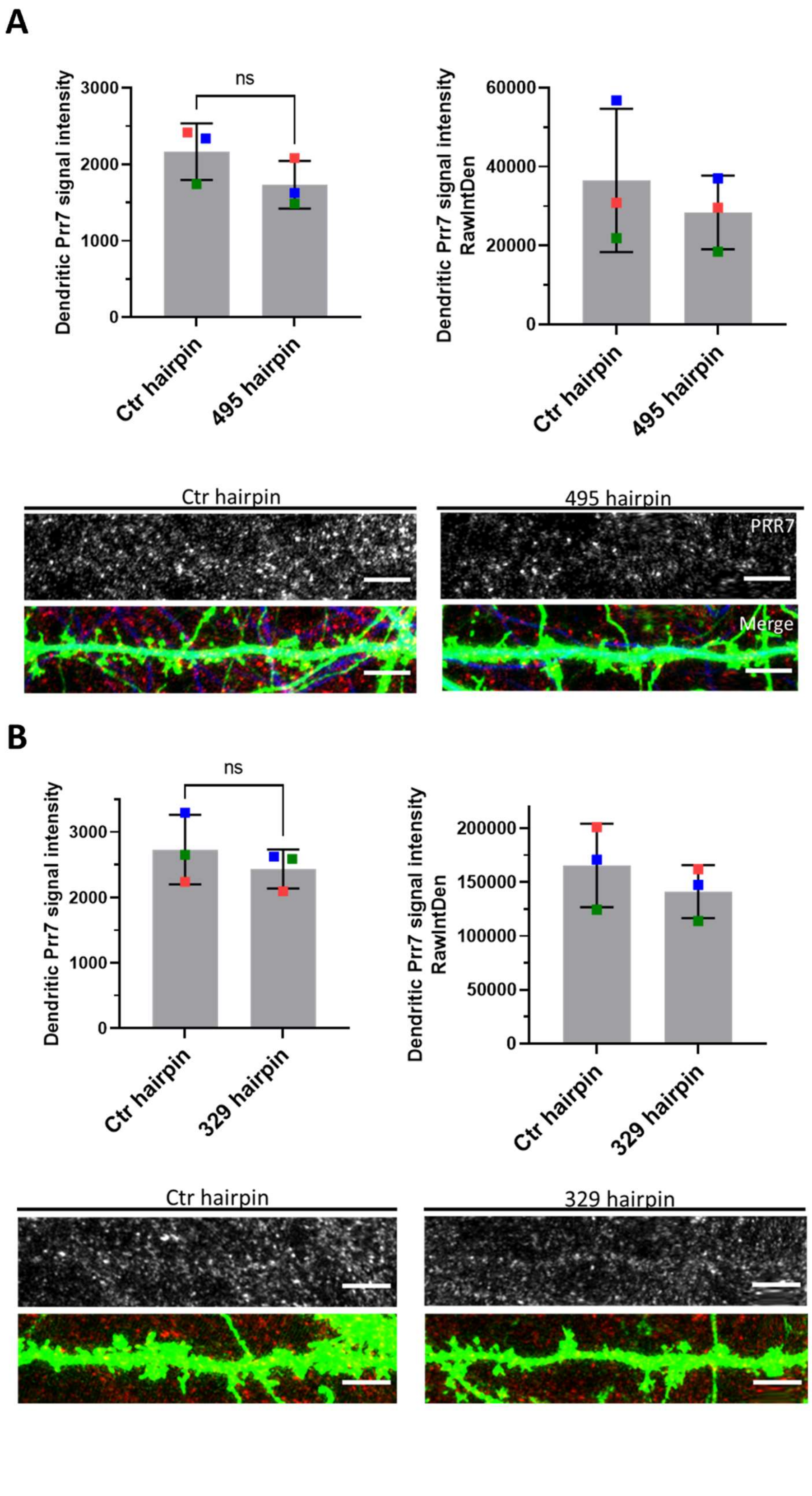


### Supplemental Figure 9.

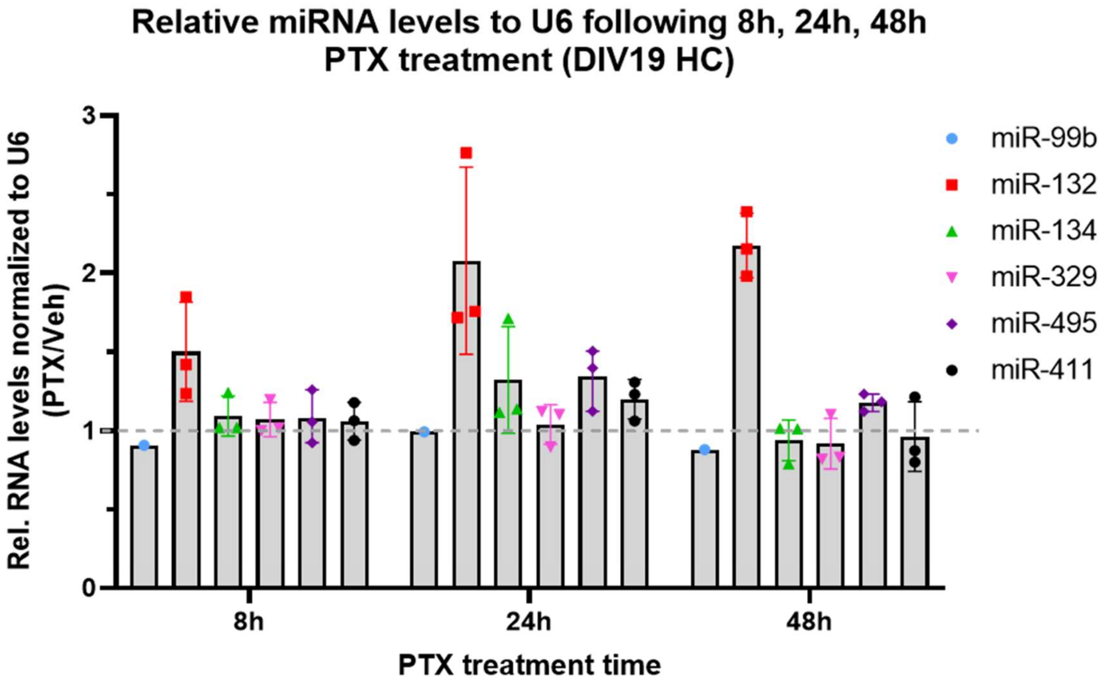


**Supplemental Figure 10.**

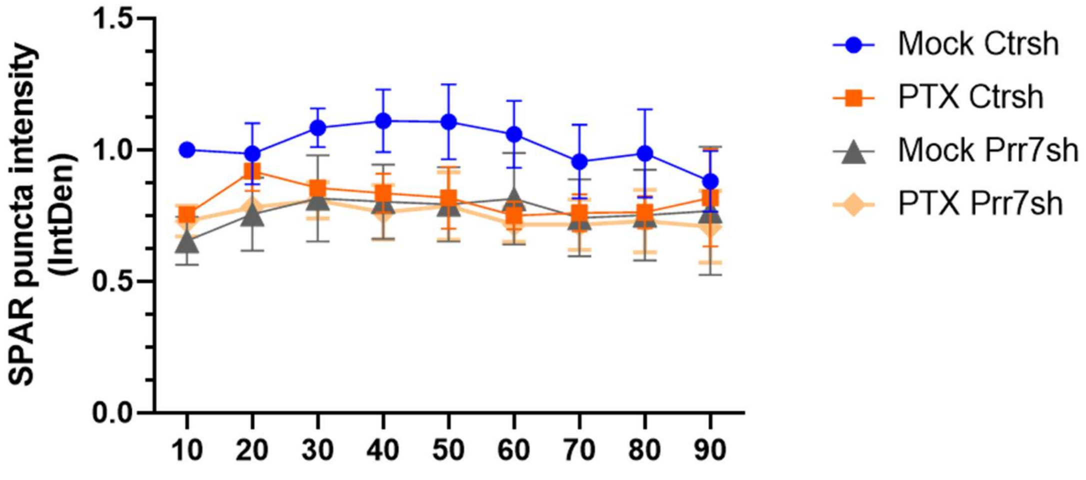


**Supplemental Figure 11.**

#### Supplemental Data

GluA1 and SPAR western blots- all replicates

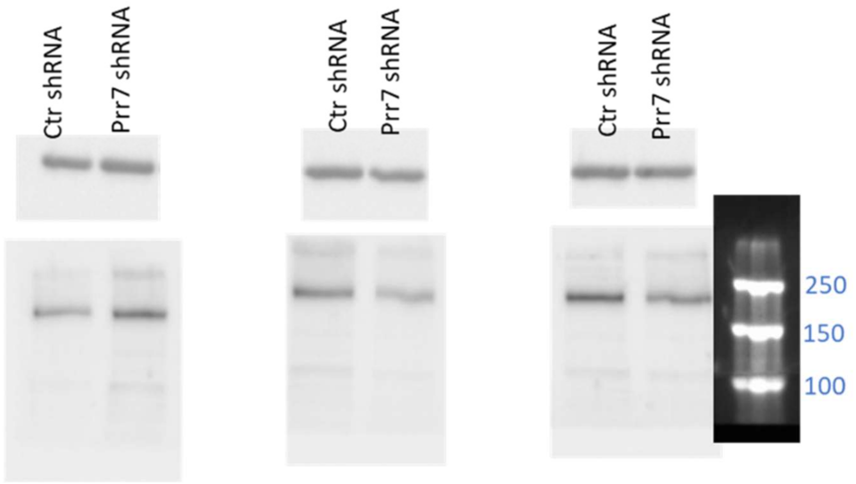


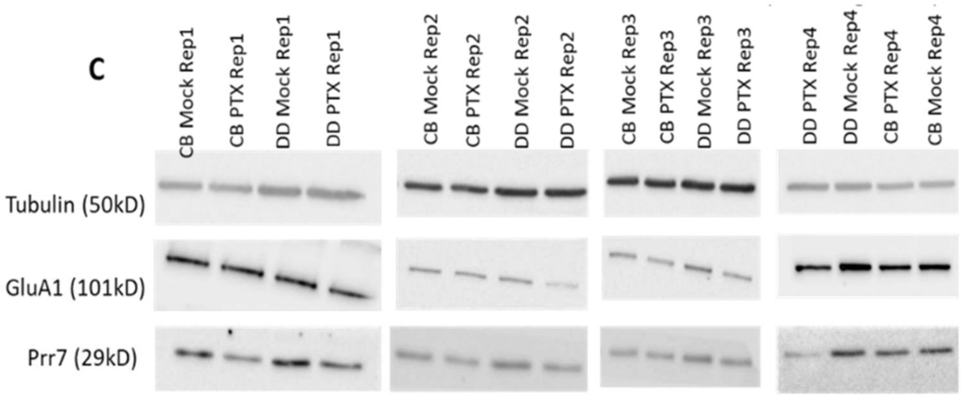


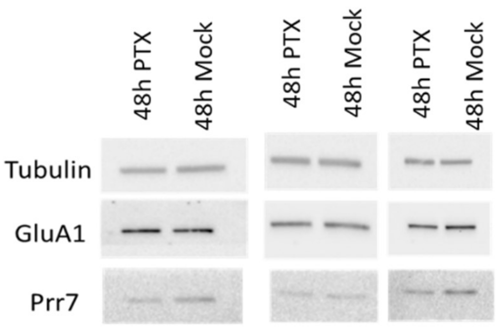

### Cell segments used for dendrite closeups

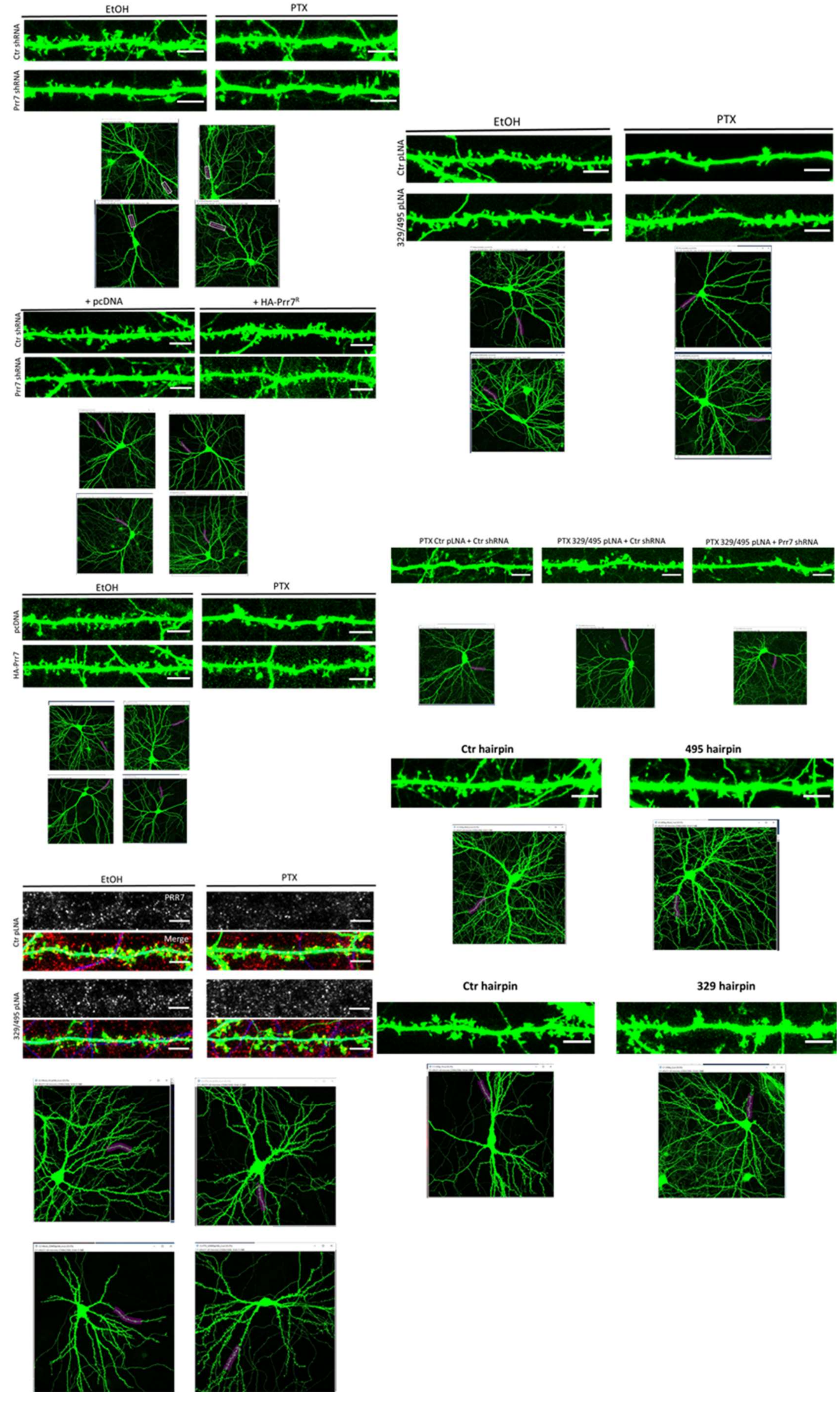

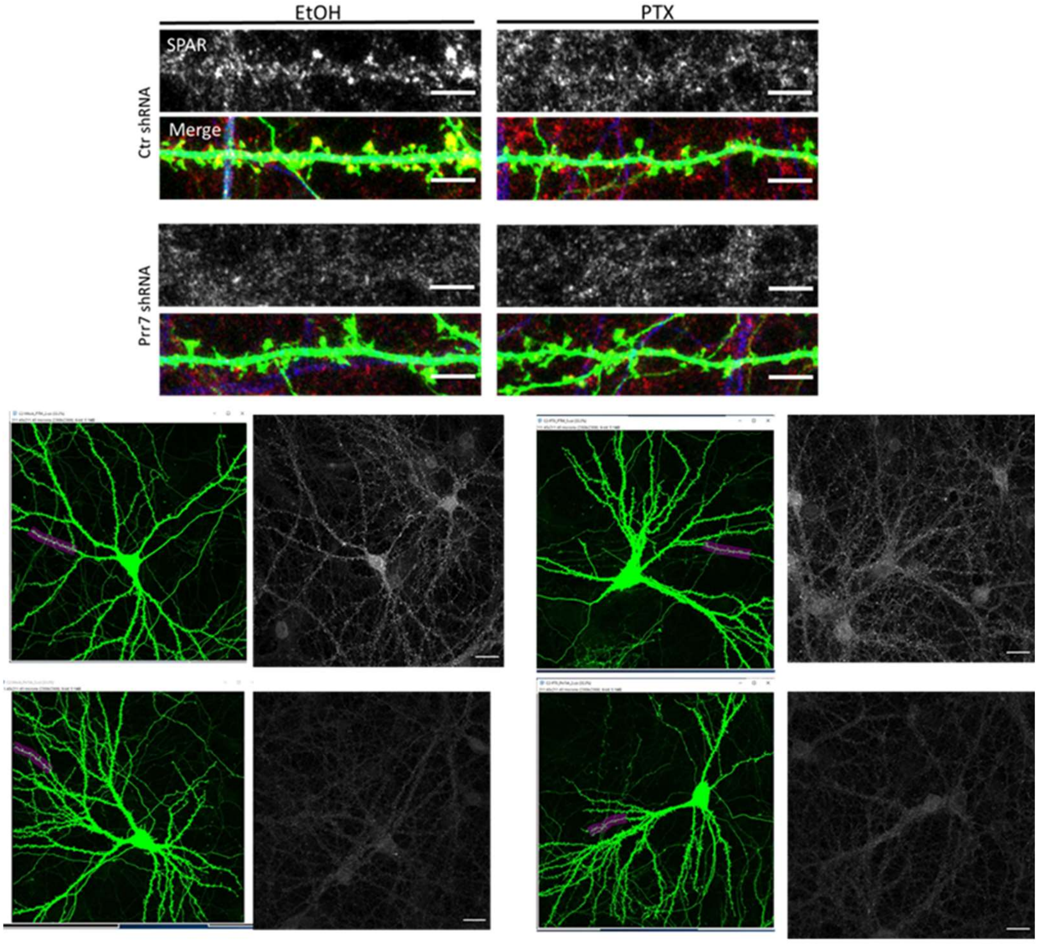
EtOH
PTX
SPAR
Merge
Ctr shRNA
Prr7 shRNA

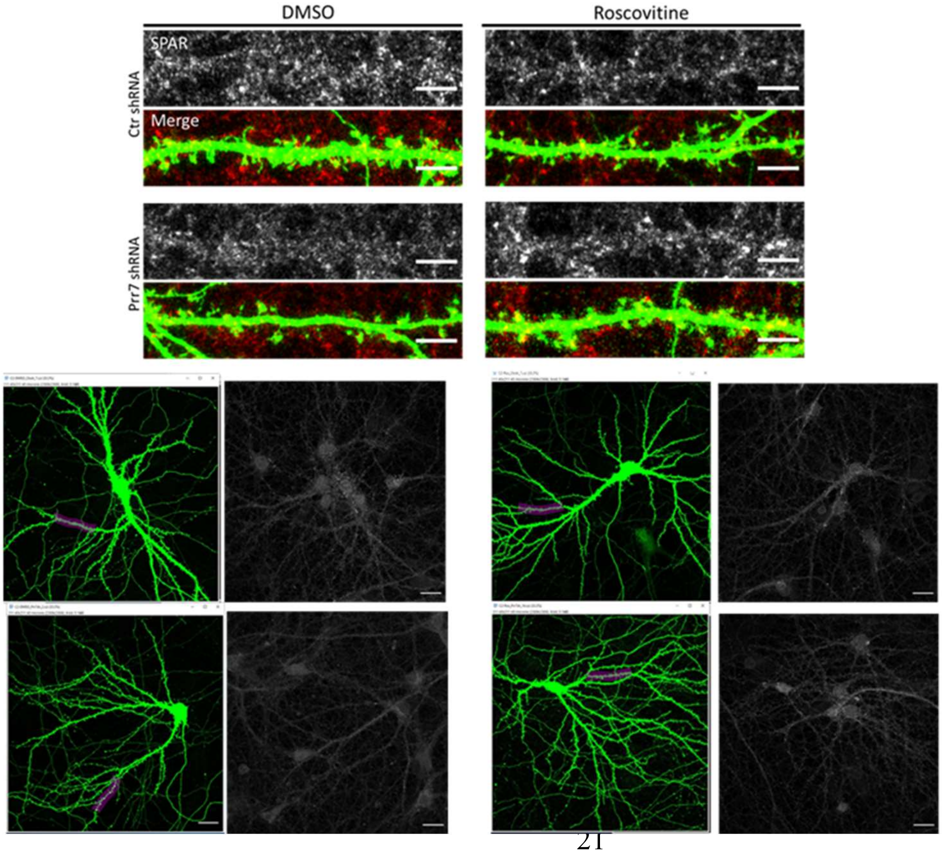
DMSO
Roscovitine
SPAR
Merge
Ctr shRNA
Prr7 shRNA
